# Chromosomal rearrangements and sequence similarity drive preferential allosyndetic introgression from a wild relative into wheat

**DOI:** 10.64898/2026.03.31.715614

**Authors:** Honglian Ye, Qijun Zhang, Prakitchai Chotewutmontri, Swarupa Nanda Mandal, Zhixia Niu, Yunming Long, Jianqiang Shen, Rebecca B. Whetten, Genqiao Li, Yue Jin, Sam Gale, Timothy L. Friesen, Amanda Peters Haugrud, Xiangyang Xu, Justin D. Faris, Shengming Yang, Christina Cowger, Jianli Chen, Xiwen Cai, Xiaofei Zhang, Sheng Luan, Yong Q. Gu, Daryl L. Klindworth, Steven S. Xu

## Abstract

Recombination in polyploid genomes is generally constrained to homologous or homoeologous chromosomes; however, how chromosomal rearrangements influence recombination between chromosomes remains unclear. Here, we demonstrate that large-scale chromosomal rearrangements in the wild relatives of wheat are associated with recombination involving non-homoeologous chromosomes or arms during alien gene introgression under conditions that permit homoeologous recombination mediated by *ph1b*. Using a wheat chromosome 6A monosomic-induced 6AS•6CL Robertsonian translocation combined with *ph1b*-mediated recombination, we generated 17 independent recombinants carrying a new stem rust resistance gene, *Sr69*, from *Aegilops caudata* chromosome arm 6CL. Unexpectedly, 94.1% (16 of 17) of recombinants resulted from exchanges with wheat group-7 chromosomes rather than with the homoeologous group-6 chromosome. Comparative sequence- and marker-based analyses identified a 67-Mb rearranged interval on *Ae. caudata* 6CL that corresponds to telomeric regions of the long arms of wheat group-7 chromosomes. Sequence similarity within this interval was quantitatively associated with recombination frequency, with higher similarity corresponding to more frequent translocations. Physical and optical mapping showed that recombination within the rearranged interval generated compensating 7A/6C, 7B/6C, and 7D/6C translocations, whereas recombination outside this region produced non-compensating 6A/6C exchanges. An independent case involving the powdery mildew resistance gene *Pm7C* showed a similar correspondence between a rearranged 7CL region and preferential introgression into wheat 7DS. Together, these results indicate that *ph1b*-mediated recombination involving structurally altered chromosomes is driven by local chromosomal structure and sequence similarity rather than strict homoeologous group identity. This provides a mechanistic basis for harnessing untapped beneficial genes from structurally rearranged alien genomes.

**Significance Statement:** Alien gene introgression is a powerful strategy for wheat improvement, typically relying on *ph1b*-mediated recombination between homoeologous chromosomes. The genomic basis and outcomes of introgression from structurally rearranged alien chromosomes remain unclear. Here, we show that *ph1b*-induced recombination can efficiently target wheat-allosyntenic blocks in rearranged alien genomes, preferentially transferring genes from structurally altered alien segments into their syntenic regions on wheat chromosomes of different homoeologous groups. Crossover formation is governed by extended sequence similarity within corresponding intervals rather than strict collinearity across entire homoeologous chromosomes. As many wild species exhibit extensive genome rearrangement, these findings and methodologies expand access to underexploited genetic diversity embedded within highly rearranged wild genomes for wheat improvement.

## Introduction

Wheat (*Triticum aestivum* L.; 2*n* = 6*x* = 42, AABBDD) is one of the world’s most important food crops, providing essential nutrition and serving as a major source of feed and energy (1). However, domestication bottlenecks and intensive breeding have substantially reduced its genetic diversity in modern cultivars, increasing their vulnerability to emerging biotic and abiotic stresses (2). Wild relatives within the tribe Triticeae, represent a major reservoir of genetic variation, and numerous beneficial genes have been introgressed into wheat via chromosome translocations, including the widely deployed wheat–rye 1B/1R and wheat–*Aegilops ventricosa* 2A/2N translocations (3–5).

Introgression from wild relatives in wheat has largely relied on inducing meiotic pairing and recombination between homoeologous chromosomes. The *Ph1* (pairing homoeologous 1) locus on chromosome 5B ensures homologous synapsis (autosyndesis) while suppressing pairing between homoeologous chromosomes (allosyndesis) (6, 7). *Ph1* spans a complex ∼2.5-Mb region containing subtelomeric heterochromatin and *cdc2*-related genes (8). In the absence of *Ph1*, as in *ph1b* or *ph1c* mutants, homoeologous recombination is permitted and has been widely used in pre-breeding to transfer genes from wild species into wheat (9–12). Most *ph1b*-mediated introgression efforts have focused on donor species with chromosomes broadly colinear with wheat, because collinearity facilitates homoeologous pairing and formation of stable translocations.

In contrast, many Triticeae species harbor extensive chromosomal rearrangements that disrupt homoeologous collinearity with the wheat genome, including rye (13), *Ae. caudata* (14, 15), *Ae. umbellulata* (16, 17), and *Ae. uniaristata* (18). Although these species represent rich sources of genetic variation, their structural divergence raises two fundamental questions regarding recombination: Can genes within rearranged chromosomal segments be efficiently transferred into wheat, and what factors determine recombination patterns when homoeologous pairing is permitted (e.g., in *ph1b* mutants)? Thus, the mechanisms underlying recombination among structurally complex chromosomes remain a central and unresolved challenge in polyploid genome biology.

Among structurally divergent wheat relatives, *Ae. caudata* (2n = 2x = 14, CC) is a cytogenetically well characterized system for examining the effects of structural rearrangement. Six disomic addition (DA) lines, each carrying individual *Ae. caudata* chromosomes from accession S740-69 in the wheat cultivar ‘Alcedo’, were previously developed (19, 20). Of these, five lines contain *Ae. caudata* chromosomes with translocations or inversions relative to wheat (14, 15, 21). Notably, the *Ae. caudata* chromosomes in DA lines AIII(D) and AIV(F) were assigned to 6C and 7C, respectively, but exhibit interchromosomal and intrachromosomal rearrangements. The telomeric region of the 6C long arm (6CL) is syntenic with the telomeric regions of the long arms of wheat group-7 chromosomes (group-7L), whereas the telomeric region of 7CL corresponds to the regions of the short arms of wheat group-7 chromosomes (group-7S) (14, 15). Diseases assays revealed that AIII(D) and AIV(F) confer resistance to multiple isolates of powdery mildew pathogen *Blumeria graminis* f. sp. *tritici* (*Bgt*), and that AIII(D) also conferred resistance to diverse races of stem rust pathogen *Puccinia graminis* f. sp. *tritici* (*Pgt*) (21), making them valuable resources for studying gene introgression from structurally complex genomes.

Here, we used a chromosome engineering system to introgress a stem rust resistance (*Sr*) gene from *Aegilops caudata* chromosome 6C into wheat under p*h1b*-mediated recombination. This system provides an opportunity to examine how recombination occurs between structurally rearranged chromosomal segments in a polyploid genome. We examined whether genes embedded in rearranged chromosomal regions can be efficiently transferred into wheat and what factors influence recombination when homoeologous paring constraints are relaxed. Notably, the powdery mildew resistance gene *Pm7C* was recently transferred from AIV(F) chromosome arm 7CL to wheat chromosome 7DS (22), suggesting that genetic exchange can occur between chromosomal regions that do not follow classical homoeologous group relationships. Building on this observation, we investigate how chromosomal structure and sequence relationships shape recombination and gene transfer between the short (p) and long (q) arms of homoeologous chromosomes.

## Results

### Introgression of the *Sr* gene reveals unexpected recombination patterns

To introgress the *Sr* gene from *Ae. caudata* into wheat, we generated a population of 223 F₂ plants from double monosomic F₁ plants (20″ + 1′_6A_ + 1′_6C_), derived from a cross between Chinese Spring (CS) monosomic 6A (M6A) and AIII(D). The population was segregated into 134 susceptible and 89 resistant plants when screened with *Pgt* race TMLKC (*SI Appendix*, *Method S1*, Fig. S1 and Table S1). Genomic *in situ* hybridization (GISH) analysis of 85 resistant F₂ plants showed that 50 (58.8%) carried one alien chromosome, 29 (34.1%) carried two, five (5.9%) carried one alien plus one translocated chromosome, but only one plant (1.2%; 12N61-16) possessed a 6AS•6CL Robertsonian translocation (*SI Appendix*, Fig. S2*C*). This plant (12N61-16) was crossed with the CS *ph1b* mutant. Resistant progenies were backcrossed to CS *ph1b*, and 74 BC₁F₁ plants were screened for stem rust resistance and *ph1b* homozygosity. Resistant *ph1b*-homozygous BC₁F₁ plants were backcrossed to CS to produce a BC₂F₁ population for recombinant selection.

Among 1,039 BC₂F₁ plants evaluated with TMLKC, 471 were resistant and 568 susceptible, deviating from the expected 1:1 ratio (χ² = 9.06, *P* = 0.003). All resistant plants and 245 randomly selected susceptible plants were genotyped with seven SSR markers (*Wmc232*, *Wmc621*, *Wmc773*, *Dupw217*, *Cfd6*, *Gdm147*, and *Gwm332*) linked to the *Ae. caudata* 6CL arm (Tables 1 and *SI Appendix*, Table S2). Marker dissociation occurred in 27 plants (17 resistant and 10 susceptible), establishing marker order (except *Gdm147* and *Wmc773*) with *Cfd6* and *Wmc232* as the most proximal and distal loci, respectively. All resistant recombinants retained the *Ae. caudata* allele at *Wmc232*, whereas all susceptible recombinants lacked it. Notably, 15 of 17 resistant recombinants (88.2%) retained only *Wmc232*, while 6 of 10 susceptible recombinants (60%) retained all *Ae. caudata* markers except *Wmc232*, indicating a tight linkage between the *Sr* gene and the distal region (*Wmc232*). These marker patterns revealed an unexpected recombination outcome, in which crossover breakpoints were strongly concentrated in the distal region of 6CL rather than distributed across homoeologous chromosome arms.

GISH analysis confirmed that all 17 resistant recombinants retained *Ae. caudata* chromatin at the telomeric region of the translocated arm (*SI Appendix*, Figs. S2–S4). In contrast, susceptible recombinant 6-0303 lacked the *Ae. caudata* allele at *Wmc232* and instead carried a pair of translocation chromosomes with a small wheat telomeric segment (*SI Appendix*, Fig. S2I). Another susceptible line, 6-0142, possessed chromosomes indistinguishable from the 6AS•6CL Robertsonian translocation (*SI Appendix*, Fig. S3*C*); this suggests that the missing *Ae. caudata* segment carrying *Wmc232* and the *Sr* gene, was too small for GISH detection. We therefore hypothesized that these unexpected recombination patterns reflect underlying structural features of the *Ae. caudata* 6CL.

### A rearranged distal 6CL region is associated with wheat group 7 chromosomes

Marker analysis revealed a key structural feature of the *Ae. caudata* chromosome from AIII(D). While five markers (*Cfd6*, *Gdm147*, *Wmc773*, *Dupw217*, and *Wmc621*) mapped to wheat homoeologous group 6, the two most distal markers (*Gwm332* and *Wmc232*) mapped to group-7 chromosomes (Table 1). This discordance indicates that the distal 6CL region is syntenic with wheat group-7 chromosomes. Breakpoint analysis supported this conclusion: 15 of 17 resistant recombinants carried the wheat allele at *Gwm332* but retained the *Ae. caudata* allele at *Wmc232*, placing crossovers within the distal group-7L syntenic segment. One recombinant (6–0241) exhibited a breakpoint in the proximal group-6 region. These results demonstrate that the *Sr* gene resides in the distal 6CL segment that aligns with wheat group-7 chromosomes rather than group-6 chromosomes.

**Table 1.** Presence of *Aegilops caudata* chromatin in BC2F1 plants carrying allosyndetic recombinants, as determined using seven wheat SSR markers.

| Plant No. <sup>a</sup> | Reaction to Pgt race TMLKC | Presence of wheat (W) and *Ae. caudata* (C) chromatins detected by markers<sup>b</sup> |  |  |  |  |  |  | No. of Plants |
| --- | --- | --- | --- | --- | --- | --- | --- | --- | --- |
|  |  | *Cfd6*<br>(7A/3A/6A) | *Gdm147*<br>(6A/6B) | *Wmc773*<br>(5B/6D) | *Wmc621*<br>(6A/6B) | *Dupw217*<br>(6B) | *Gwm332*<br>(7A) | *Wmc232*<br>(4A/7B) |  |
| 6-0015 | R | W | W | W | W (null) | W | W (null) | C | 15<sup>c</sup> |
| 6-0809 | R | W | W | W | W (null) | W | C | C | 1 |
| 6-0241 | R | W | W | W | C | C | C | C | 1 |
| 6-0219 | S | C | W | W | W (null) | W | W (null) | W (null) | 3<sup>d</sup> |
| 6-0144 | S | C | C | C | C | W | W (null) | W (null) | 1 |
| 6-0142 | S | C | C | C | C | C | C | W (null) | 6<sup>e</sup> |

<sup>a</sup> BC<sub>2</sub>F<sub>1</sub> plants have the pedigree of CS/4/CS *ph1b*/3/CS M6A 6A/AIII(D)//CS *ph1b*. They were generated by first backcrossing an F<sub>2</sub> plant (CS M6A 6A/AIII(D)) carrying a 6AS•6CL Robertsonian translocation to the CS *ph1b* mutant, followed by a second backcross to CS.
<sup>b</sup> (21, 30–32), GrainGenes (https://graingenes.org/cgi-bin/GG3/browse.cgi?class=marker), or were determined in this study.
<sup>c</sup> Other plants include 6-0034, 0111, 0185, 0244, 0281, 0283, 0480, 0676, 0724, 0725, 0806, 0848, 0942, and 1038.
<sup>d</sup> Other plants include 6-0351 and 0482.
<sup>e</sup> Other plants include 6-0156, 0218, 0303, 0369, and 0371.

Because *Wmc232* is dominant, we screened additional 14 wheat group-6 and 20 group-7 markers and identified a single co-dominant marker, *BF145935* (Fig. 1A and *SI Appendix*, Table S3) (23, 24). Genotyping of BC₂F₂ populations from the 17 recombinants revealed that the *Ae. caudata* 6CL segments had translocated to multiple wheat chromosomes (Figs. 1A and *SI Appendix*, Table S5). Fifteen lines carried either 7A/6C (12 lines) or 7D/6C (three lines) translocations, whereas two lines (6-0241 and 6-0244) carrying both 7A and 7D fragments, could not be genotyped with *BF145935*.

**Figure 1.**
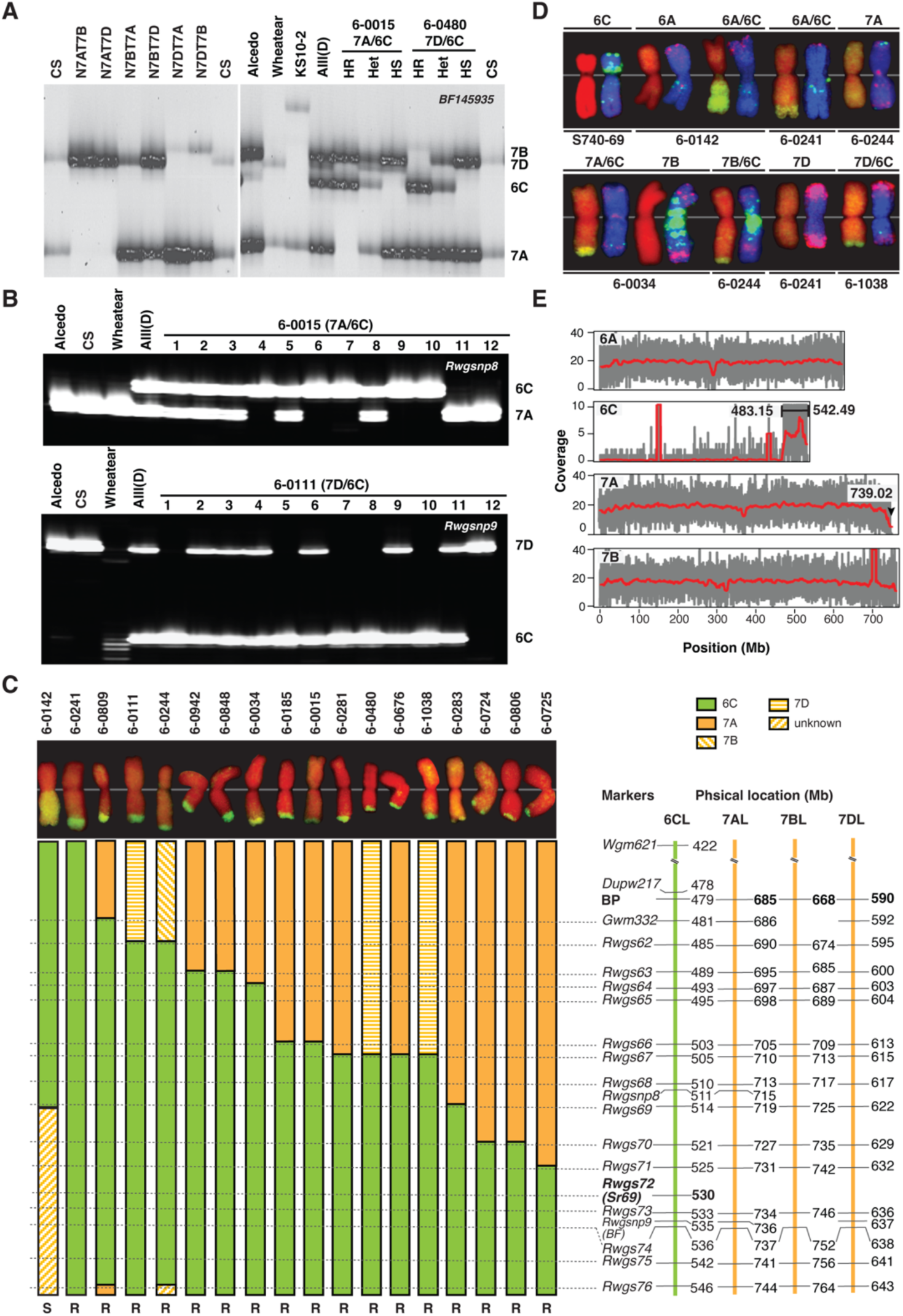
Identification and physical localization of translocation chromosomes in wheat–*Ae. caudata* introgression lines. **A**. BC_2_F_2_ populations derived from 17 recombinant BC_2_F_1_ plants with marker *BF145935* to determine translocation identity. Gel images show bands in CS nullisomic–tetrasomic lines (left), AIII(D), homozygous resistant (HR), heterozygous (Het), and susceptible (HS) plants from recombinants 6-0015 and 6-0480, with checks Wheatear (*Sr25*) and KS10-2 (*Sr43*). Marker BF145935 is diagnostic for *Th. ponticum* segments carrying Sr25 on chromosome 7D. Lines 6-0015 and 6-0480 carry 7A/6C and 7D/6C translocations, respectively (all recombinants in Fig. S5). **B**. Segregation in BC_2_F_2_ populations of 6-0015 (7A/6C) and 6-0111 (7D/6C) using STARP markers *Rwgsnp8* and *Rwgsnp9*. Populations were phenotyped with *Pgt* race TMLKC prior to DNA extraction. Testing of ≥25 BC₂F₃ progeny per resistant BC₂F₂ plant confirmed homozygous resistant 7A/6C individuals in 6-0015 (#4, #6, #7, #9, #10) and homozygous resistant 7D/6C individuals in 6-0111 (#1, #5, #7, #8, #10). Genotype cluster plots are shown in Fig. S6. **C.** Distribution of 6CL segments across 17 *Sr69* introgression lines and the susceptible line 6-0142. The top panel shows representative GISH images of the translocation chromosomes in each line. The linkage map (right) indicates the physical positions (Mb) of markers in the 6CL telomeric region of the *Ae. caudata* genome Aecau_v1 (15). BP denotes the breakpoint at which the rearranged *Ae. caudata* 6CL region begins. **D**. GISH and oligo-FISH identification of translocations in lines 6-0142, 6-0241, and 6-0244. Each chromosome is shown by GISH (left) and oligo-FISH (right), except for *Ae. caudata* chromosome 6C, which was DAPI-stained (left image). Whole-cell spreads are in Figs. S3 and S4. Lines 6-0142 and 6-0241 carry 6A/6C translocations, whereas 6-0244 carries a 7B/6C translocation. The images of 6A and 6A/6C in 6-0142, 7A and 7B/6C in 6-0244, and 7B and 7A/6C in 6-0034 were extracted from the same cells, respectively. **E.** Bionano optical mapping of line 6-0244 showing alignment coverage against the *Ae. caudata* Aecau_v1 and CS IWGSC RefSeq v2.1 (25) genomes. A 59.3-Mb *Ae. caudata* segment was mapped to the distal region (483.15–542.49 Mb) of chromosome arm 6CL, with no significant deletion detected on chromosome 7B, indicating that 6-0244 is a non-reciprocal 7B/6C translocation line. Notably, a 5-Mb telomeric region is missing from wheat chromosome arm 7AL.

To further resolve these 17 BC₂F₂ lines, we developed two co-dominant STARP (semi-thermal asymmetric reverse PCR) markers, *Rwgsnp8* and *Rwgsnp9* (*SI Appendix*, Table S3; Figs. 1B and S6). While these markers clearly distinguished 7A/6C and 7D/6C translocations, they still failed to resolve lines 6-0241 and 6-0244. Oligonucleotide multiplex fluorescence *in situ* hybridization (oligo-FISH) was then used to analyze these translocation chromosomes, using representative 7A/6, 7D/6C, and susceptible 6A/6C (line 6-0142) introgressions as controls. These analyses confirmed that line 6-0241 carries a 6A/6C translocation, while 6-0244 carries a 7B/6C translocation (Fig. 1D and *SI Appendix*, Figs. S3, and S4). These results demonstrate that recombinants predominantly involve wheat group-7 chromosomes, suggesting that recombination is not constrained by classical homoeologous group identity.

### Recombination within a rearranged distal 6CL region generates compensating translocations

The *Ae. caudata* S740-69 assembly Aecau_v1 (15) showed that the *Sr* gene, designated *Sr69*, was located at 530 Mb in the 6CL telomeric region, with a ∼67 Mb rearranged interval (479–546 Mb) that is syntenic with wheat group-7L telomeric regions (*SI Appendix*, Table S4).

To define introgression boundaries, we developed 15 PCR markers (*Rwgs62*–*Rwgs76*) (*SI Appendix*, Table S5) across the 485–546 Mb region at ∼5-Mb resolution based on conserved genes syntenic between Aecau_v1 and CS IWGSC RefSeq v2.1 (25). Except for *Rwgs72*, derived from the *Sr69* candidate gene (*Aecau6C01G127270*), the remaining markers amplified polymorphic fragments from syntenic regions of 6CL and wheat 7AL, 7BL, and 7DL (Figs. 2A and S7; Table S6). These markers enabled estimation of introgressed 6CL segments and the corresponding replaced wheat segments. For the two 6A/6C lines (6-0142 and 6-0241), four SSR markers (*Wmc621*, *Dupw217*, *Gwm332*, and *Wmc232*) further extended estimates of the gained 6CL and replaced 6AL segments.

**Figure 2.**
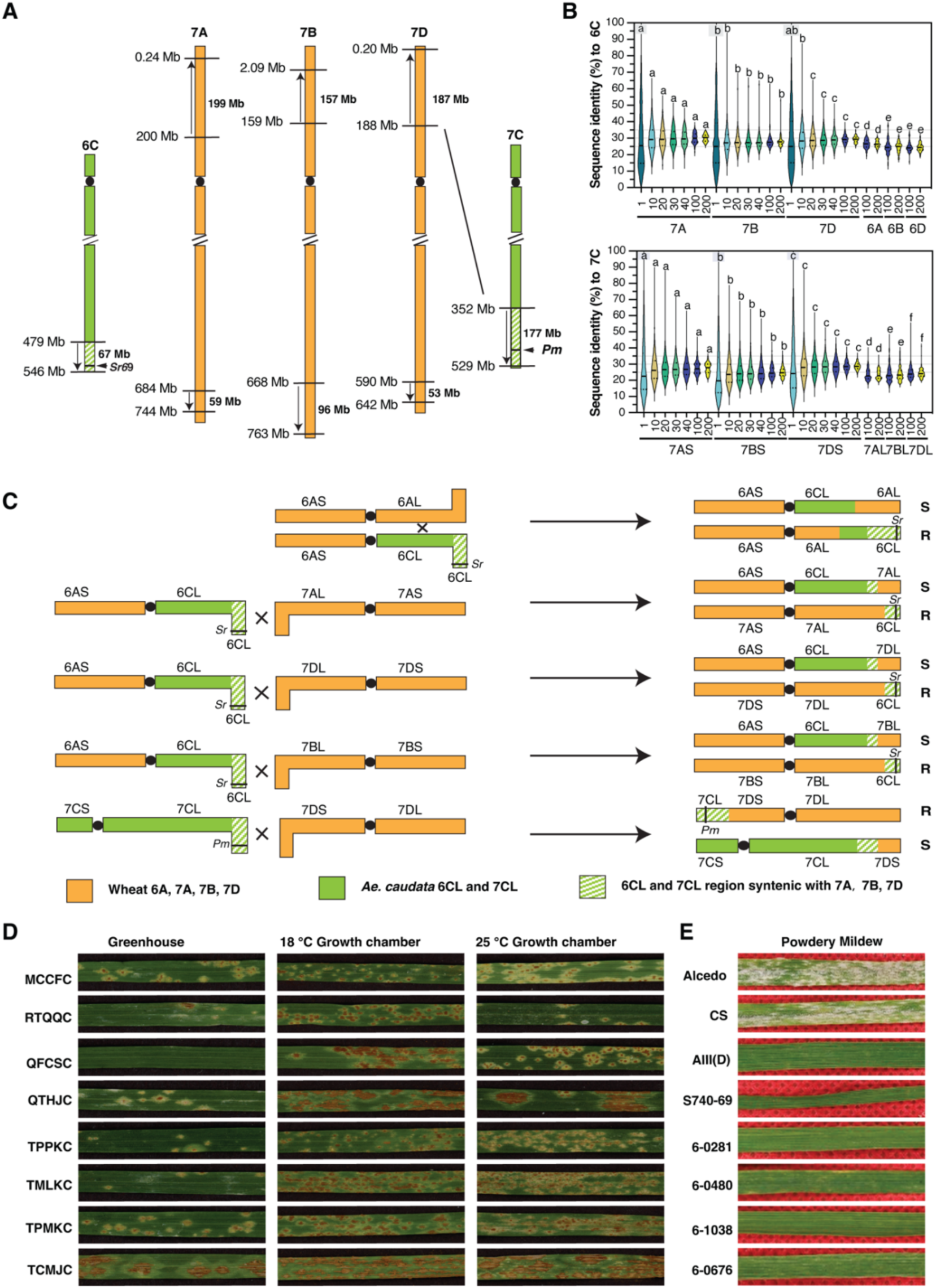
Syntenic relationship of the Ae. caudata rearranged the 6CL and 7CL regions with wheat group-7 chromosomes, allosyndetic recombination, and disease responses of wheat–Ae. caudata introgression lines. **A.** Diagram showing the physical locations of representative polymorphic markers between *Ae. caudata* chromosome arm 6CL, 7CL and wheat group 7 chromosomes. The rearranged 6CL region (67 Mb) containing *Sr69* is syntenic with the telomeric regions of chromosome arms 7AL (59 Mb), 7BL (96 Mb), and 7DL (53 Mb) based on the *Ae. caudata* Aecau_v1 (15) and CS IWGSC RefSeq v2.1 (25) genomes (Table S4). The 177-Mb distal 7CL region is syntenic with telomeric regions of wheat 7AS (199 Mb), 7BS (157 Mb), and 7DL (187 Mb). **B.** Percent sequence identity between the 6CL(top) and 7CL(buttom) syntenic region and each corresponding group-7 chromosome segment. Different letters above the violin plots indicate significant differences (adjusted *p* <0.0001) among chromosome groups within each window size; groups sharing the same letter are not significantly different. **C.** Diagrams illustrating possible modes of allosyndetic recombination involving the Robertsonian translocation chromosome 6AS•6CL with chromosomes 6A, 7A, 7D, and 7B (upper four). The bottom row illustrates recombination of 7CS•7CL with chromosome 7D, producing 7CL–7DS•7DL and 7CS•7CL–7DS recombinants for Pm7C introgression(22). Crossovers outside the rearranged 6CL region produce 6A/6C translocations, such as susceptible (S) 6AS•6CL-6AL and resistant (R) 6AS•6AL-6CL recombinants. Recombination within the rearranged 6CL region generates 7A/6C, 7B/6C, and 7D/6C translocations, depending on pairing of the rearranged 6CL region with its homoeologous regions on chromosomes 7A, 7D, and 7B, respectively. **D.** Reactions of the wheat–*Ae. caudata* 7A/6C introgression line 6-0015 to eight *Puccinia graminis* f. sp. *tritici* races (left) evaluated under greenhouse and growth chamber conditions (18 °C and 25 °C). **E.** Reactions of wheat cultivars ‘Aceldo’ and ‘Chinese Spring’, *Ae. caudata* accession S740-69, the Aceldo–*Ae. caudata* S740-69 disomic addition line AIII(D), and four resistant CS–*Ae. caudata* 7A/6C and 7D/6C translocation lines when tested with *Blumeria graminis* f. sp. *tritici* (Bgt) isolate OKS(14)-B-3-1.

Most *Sr69* introgression lines (14 of 17) carried compensating terminal translocations, in which the distal 6CL replaced terminal regions of wheat chromosomes (Table 2). Specifically, 12 lines possessed 7A/6C translocations, including one intercalary configuration (line 6-0809, 7AS•7AL-6CL-7AL), and 11 terminal configurations (7AS•7AL-6CL; *SI Appendix*, Table S6). Additionally, three lines (6-0111, 6-0480, and 6-1038) carried compensating 7DS•7DL-6CL translocations, and one line (6–0244) carried a 7BS•7BL-6CL translocation. In contrast, line 6-0241 harbored a 6AS•6AL-6CL translocation.

**Table 2.**
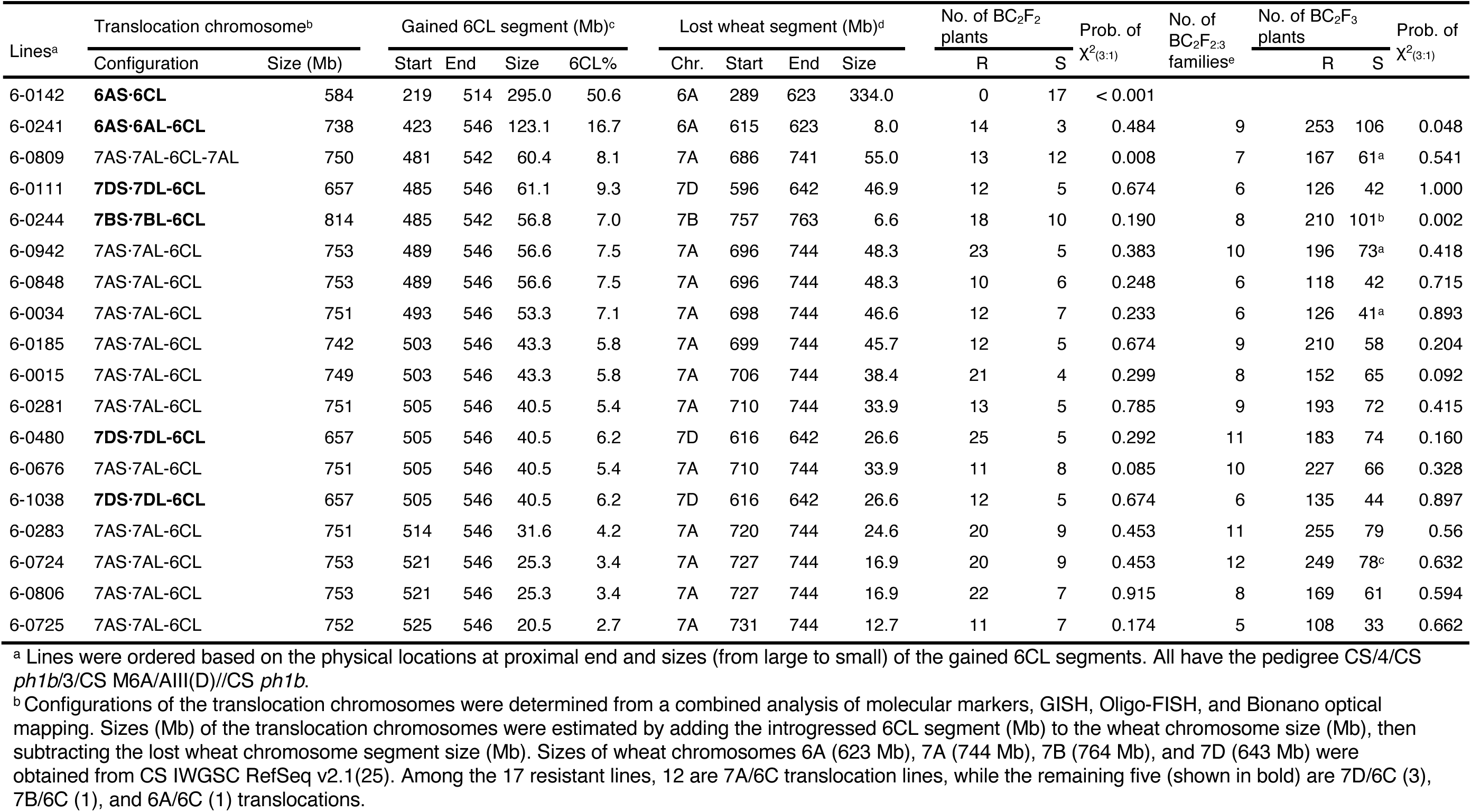
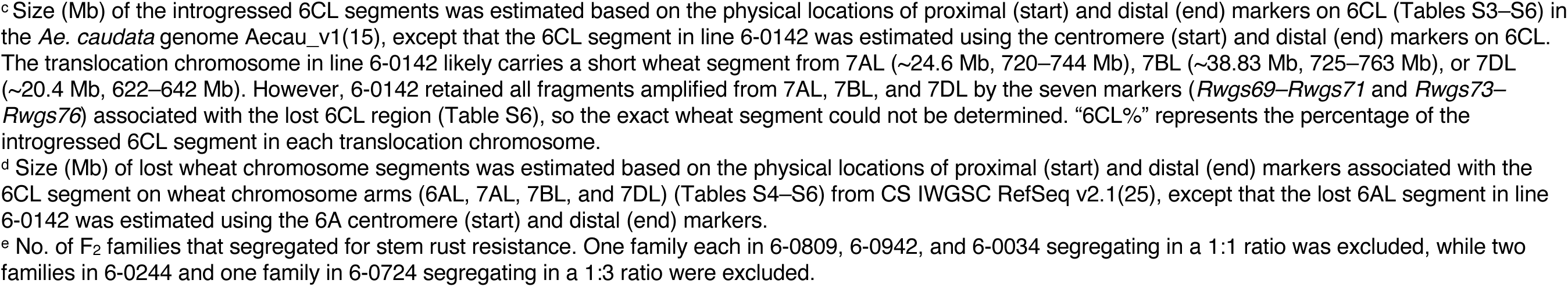
Identified translocation chromosomes, estimated locations and sizes of gained 6CL segments, lost wheat chromosome segments in 17 wheat–*Ae. caudata Sr69* introgression lines, and segregation analysis of stem rust resistance in BC2F2 populations and BC2F2:3 families.

| Lines<sup>a</sup> | Translocation chromosome<sup>b</sup> | | Gained 6CL segment (Mb)<sup>c</sup> | | | | Lost wheat segment (Mb)<sup>d</sup> | | | | No. of BC<sub>2</sub>F<sub>2</sub> plants | | Prob. of $\chi^2_{(3:1)}$ | No. of BC<sub>2</sub>F<sub>2:3</sub> families<sup>e</sup> | No. of BC<sub>2</sub>F<sub>3</sub> plants | | Prob. of $\chi^2_{(3:1)}$ |
| --- | --- | --- | --- | --- | --- | --- | --- | --- | --- | --- | --- | --- | --- | --- | --- | --- | --- |
|  | Configuration | Size (Mb) | Start | End | Size | 6CL% | Chr. | Start | End | Size | R | S |  |  | R | S |  |
| 6-0142 | **6AS-6CL** | 584 | 219 | 514 | 295.0 | 50.6 | 6A | 289 | 623 | 334.0 | 0 | 17 | < 0.001 |  |  |  |  |
| 6-0241 | **6AS-6AL-6CL** | 738 | 423 | 546 | 123.1 | 16.7 | 6A | 615 | 623 | 8.0 | 14 | 3 | 0.484 | 9 | 253 | 106 | 0.048 |
| 6-0809 | 7AS-7AL-6CL-7AL | 750 | 481 | 542 | 60.4 | 8.1 | 7A | 686 | 741 | 55.0 | 13 | 12 | 0.008 | 7 | 167 | 61<sup>a</sup> | 0.541 |
| 6-0111 | **7DS-7DL-6CL** | 657 | 485 | 546 | 61.1 | 9.3 | 7D | 596 | 642 | 46.9 | 12 | 5 | 0.674 | 6 | 126 | 42 | 1.000 |
| 6-0244 | **7BS-7BL-6CL** | 814 | 485 | 542 | 56.8 | 7.0 | 7B | 757 | 763 | 6.6 | 18 | 10 | 0.190 | 8 | 210 | 101<sup>b</sup> | 0.002 |
| 6-0942 | 7AS-7AL-6CL | 753 | 489 | 546 | 56.6 | 7.5 | 7A | 696 | 744 | 48.3 | 23 | 5 | 0.383 | 10 | 196 | 73<sup>a</sup> | 0.418 |
| 6-0848 | 7AS-7AL-6CL | 753 | 489 | 546 | 56.6 | 7.5 | 7A | 696 | 744 | 48.3 | 10 | 6 | 0.248 | 6 | 118 | 42 | 0.715 |
| 6-0034 | 7AS-7AL-6CL | 751 | 493 | 546 | 53.3 | 7.1 | 7A | 698 | 744 | 46.6 | 12 | 7 | 0.233 | 6 | 126 | 41<sup>a</sup> | 0.893 |
| 6-0185 | 7AS-7AL-6CL | 742 | 503 | 546 | 43.3 | 5.8 | 7A | 699 | 744 | 45.7 | 12 | 5 | 0.674 | 9 | 210 | 58 | 0.204 |
| 6-0015 | 7AS-7AL-6CL | 749 | 503 | 546 | 43.3 | 5.8 | 7A | 706 | 744 | 38.4 | 21 | 4 | 0.299 | 8 | 152 | 65 | 0.092 |
| 6-0281 | 7AS-7AL-6CL | 751 | 505 | 546 | 40.5 | 5.4 | 7A | 710 | 744 | 33.9 | 13 | 5 | 0.785 | 9 | 193 | 72 | 0.415 |
| 6-0480 | **7DS-7DL-6CL** | 657 | 505 | 546 | 40.5 | 6.2 | 7D | 616 | 642 | 26.6 | 25 | 5 | 0.292 | 11 | 183 | 74 | 0.160 |
| 6-0676 | 7AS-7AL-6CL | 751 | 505 | 546 | 40.5 | 5.4 | 7A | 710 | 744 | 33.9 | 11 | 8 | 0.085 | 10 | 227 | 66 | 0.328 |
| 6-1038 | **7DS-7DL-6CL** | 657 | 505 | 546 | 40.5 | 6.2 | 7D | 616 | 642 | 26.6 | 12 | 5 | 0.674 | 6 | 135 | 44 | 0.897 |
| 6-0283 | 7AS-7AL-6CL | 751 | 514 | 546 | 31.6 | 4.2 | 7A | 720 | 744 | 24.6 | 20 | 9 | 0.453 | 11 | 255 | 79 | 0.56 |
| 6-0724 | 7AS-7AL-6CL | 753 | 521 | 546 | 25.3 | 3.4 | 7A | 727 | 744 | 16.9 | 20 | 9 | 0.453 | 12 | 249 | 78<sup>c</sup> | 0.632 |
| 6-0806 | 7AS-7AL-6CL | 753 | 521 | 546 | 25.3 | 3.4 | 7A | 727 | 744 | 16.9 | 22 | 7 | 0.915 | 8 | 169 | 61 | 0.594 |
| 6-0725 | 7AS-7AL-6CL | 752 | 525 | 546 | 20.5 | 2.7 | 7A | 731 | 744 | 12.7 | 11 | 7 | 0.174 | 5 | 108 | 33 | 0.662 |

<sup>a</sup> Lines were ordered based on the physical locations at proximal end and sizes (from large to small) of the gained 6CL segments. All have the pedigree CS/4/CS *ph1b*/3/CS M6A/AIII(D)//CS *ph1b*.
<sup>b</sup> Configurations of the translocation chromosomes were determined from a combined analysis of molecular markers, GISH, Oligo-FISH, and Bionano optical mapping. Sizes (Mb) of the translocation chromosomes were estimated by adding the introgressed 6CL segment (Mb) to the wheat chromosome size (Mb), then subtracting the lost wheat chromosome segment size (Mb). Sizes of wheat chromosomes 6A (623 Mb), 7A (744 Mb), 7B (764 Mb), and 7D (643 Mb) were obtained from CS IWGSC RefSeq v2.1(25). Among the 17 resistant lines, 12 are 7A/6C translocation lines, while the remaining five (shown in bold) are 7D/6C (3), 7B/6C (1), and 6A/6C (1) translocations.

Across the 17 resistant lines, proximal breakpoints ranged from 423 to 525 Mb and distal endpoints from 542 to 546 Mb. Introgressions spanned 20.5–123.1 Mb, representing 2.7–16.7% of the translocation chromosomes; the largest was found in 6-0241 and the smallest in 6-0725 (Table 2). Corresponding wheat chromatin losses ranged from 6.6 Mb on 7B (6–0244) to 55.0 Mb on 7A (6–0809). Notably, the susceptible control 6-0142 retained a large proximal 6CL segment (219–514 Mb; 295 Mb) but lacked the distal region shared by resistant lines, resulting in the largest wheat loss (334 Mb on 6A).

Gains of 6CL segments were highly correlated with losses of the corresponding wheat segments in all 7A/6C and 7D/6C lines (*r* = 0.9522, *P* < 0.0001), indicating largely reciprocal exchange. In contrast, 7B/6C line (6–0244) gained a large 56.8-Mb 6CL segment but lost only 6.6 Mb of wheat 7BL; the introgressed segment also lacked a small 4.8-Mb telomeric region corresponding to the deleted 7BL segment (Fig. 1C and *SI Appendix*, Table S6). Optical mapping independently validated these results, showing that line 6-0244 carries a 59.3-Mb 6CL terminal segment with no major 7B deletion (Fig. 1E and *SI Appendix*, Fig. S8). The absence of both 7BL and 6CL alleles at distal marker *Rwgs76* (Table S6) indicates a nonreciprocal translocation in which the large 6CL segment fused to the 7BL terminus, likely accompanied by loss of a small 7BL telomeric region. These pattens indicate that recombination outcome depends on crossover position relative to the rearranged distal 6CL region, with crossovers within this region producing compensating translocations.

Transmissions of the introgressed segments were evaluated by phenotyping stem rust resistance in the BC_2_F_2_ populations and BC_2_F_2:3_ families of 17 *Sr69* introgression lines (Table 2). Sixteen of 17 BC₂F₂ populations fit the expected 3:1 resistant-to-susceptible ratio, except line 6-0809 (13 R:12 S). In the BC_2_F_3_ generation, 147 segregating BC_2_F_2:3_ families from 17 introgression lines were analyzed. Homogeneity tests led to the exclusion of six BC_2_F_2:3_ families. The remaining families, each comprising 5–12 BC_2_F_2:3_ families, were tested for goodness of fit to a 3:1 ratio; only lines 6-0241 and 6-0244 deviated from this expectation. These results indicated normal transmission of the *Ae. caudata* segments in all the introgression lines except for line 6-0244 and possibly 6-0241.

### Sequence similarity within the rearranged region favors recombination toward group-7 chromosomes

Comparative genomic analysis showed that the 67-Mb rearranged 6CL interval aligns with the telomeric regions of wheat chromosome arms 7AL (60 Mb), 7DL (52 Mb), and 7BL (96 Mb) (15). Markers defining the proximal 6CL breakpoint mark the transition from wheat group-6 homoeology to group-7 synteny, indicating that distal 6CL forms a contiguous block aligned with group-7 arms rather than group-6 chromosomes (Fig. 2A and *SI Appendix*, Tables S2, S5 and S7). Proximal markers (*Wgm621* and *Dupw217*) retain group-6 correspondence, whereas distal markers preserve order and alignment with 7AL, 7BL, and 7DL telomeres. This structural configuration is consistent with the deviation from canonical homoeologous recombination.

To test whether this structural correspondence extends to the sequence level, we compared the rearranged 6CL region with wheat chromosomes 7A, 7B, and 7D using sliding-window analysis of aligned genomic blocks (Fig. 2B and *SI Appendix*, Fig. S9 and Table S7). At small window sizes (< 30 blocks), sequence identity distributions were broad and overlapping, indicating local heterogeneity. With increasing window size (30–200 blocks), variance decreased and chromosome-level difference became apparent. Averaging ≥100 blocks revealed significant differences among 6CL/7AL, 6CL/7BL, and 6CL/7DL (P < 0.0001) (*SI Appendix*, Table S7). At 200-block resolution, sequence identity was highest between 6CL and 7AL (30.11%), followed by 7DL (29.30%) and 7BL (28.87%) (Fig. 2B and *SI Appendix*, Table S7), whereas identities with group-6 chromosomes were consistently lower (6AL, 26.54%; 6BL, 24.71%; 6DL, 24.53%). This gradient of sequence similarity (7A > 7D > 7B) closely matched the observed translocation frequencies, suggesting that recombination is associated with higher local sequence similarity within the rearranged interval. Together with the marker-defined structure, these results provided a quantitative basis for the predominance of group-7/6C translocations.

Consistent with this model, the crossover position relative to the rearranged interval determines the recombination outcome. Crossovers outside the rearranged interval generate non-compensating 6A/6C translocations, whereas crossovers within the interval produce compensating group-7/6C exchanges (Fig. 2C). Thus, the combination of crossover positions relative to the rearranged 6CL block, chromosomal rearrangement, and sequence similarity within the distal 6CL region explains the translocation patterns in Table 2.

### An independent introgression case supports the same structural relationships

To assess whether this relationship generalizes beyond *Sr69* region, we examined an independent introgression case involving the powdery mildew resistance gene *Pm7C*, previously introgressed from *Ae. caudata* 7CL into wheat 7DS (22). The distal 177.2-Mb region of 7CL shows synteny with wheat 7AS (199.4 Mb), 7BS (156.8 Mb), and 7DS (187.4 Mb) (Fig. 2A and *SI Appendix*, Tables S4 and S8) (15). The rearranged 7CL region showed consistently higher sequence identity to its allosyntenic regions on 7DS (28.7%), 7AS (27.5%), and 7BS (25.1%) than to the corresponding regions on 7AL (23.5%), 7BL (24.0%), and 7DL (24.7%) (Fig. 2B and *SI Appendix*, Fig. S9, Tables S4 and S8). Sequence identity was highest with 7DS, consistent with preferential introgression of *Pm7C* from 7CL into 7DS (Fig. 2*C*). This independent case supports the generality of the relationship between chromosomal rearrangement, sequence similarity, and recombination outcome.

Together, these results indicate that chromosomal rearrangements create large continuous regions that share structural and sequence similarity with non-homoeologous chromosomes. Variation in sequence similarity within these regions is associated with recombination frequency, providing a mechanistic explanation for the observed introgression patterns.

### *Sr69* introgression lines show broad stem rust resistance and variable powdery mildew resistance

All 17 *Sr69* introgression lines were evaluated against 16 domestic and foreign *Pgt* races (Table 3 and *SI Appendix*, Table S1). All lines were resistant to 13 races, producing infection types (ITs) of 0;–23, indicating that *Sr69* confers broad-spectrum resistance across diverse pathogen backgrounds. Reactions to race QTHJC were variable (ITs 2–34), whereas all lines were susceptible to TCMJC and TTRTF. The 6-0142 (6A/6C) line, which lacks the terminal 6CL segment, showed high ITs (3–4) to 13 races except for JCMNC (IT 23^−^) and QCCJC (IT 2^+^), confirming that resistance in the introgression lines is associated with the distal 6CL region carrying *Sr69*. Although all 17 introgression lines were susceptible to TTRTF, both S740-69 and AIII(D) were highly resistant (IT 1^+^), indicating that chromosome 6C likely carries additional *Sr* gene(s) that were not captured in the introgressed segments (Table 3).

**Table 3.**
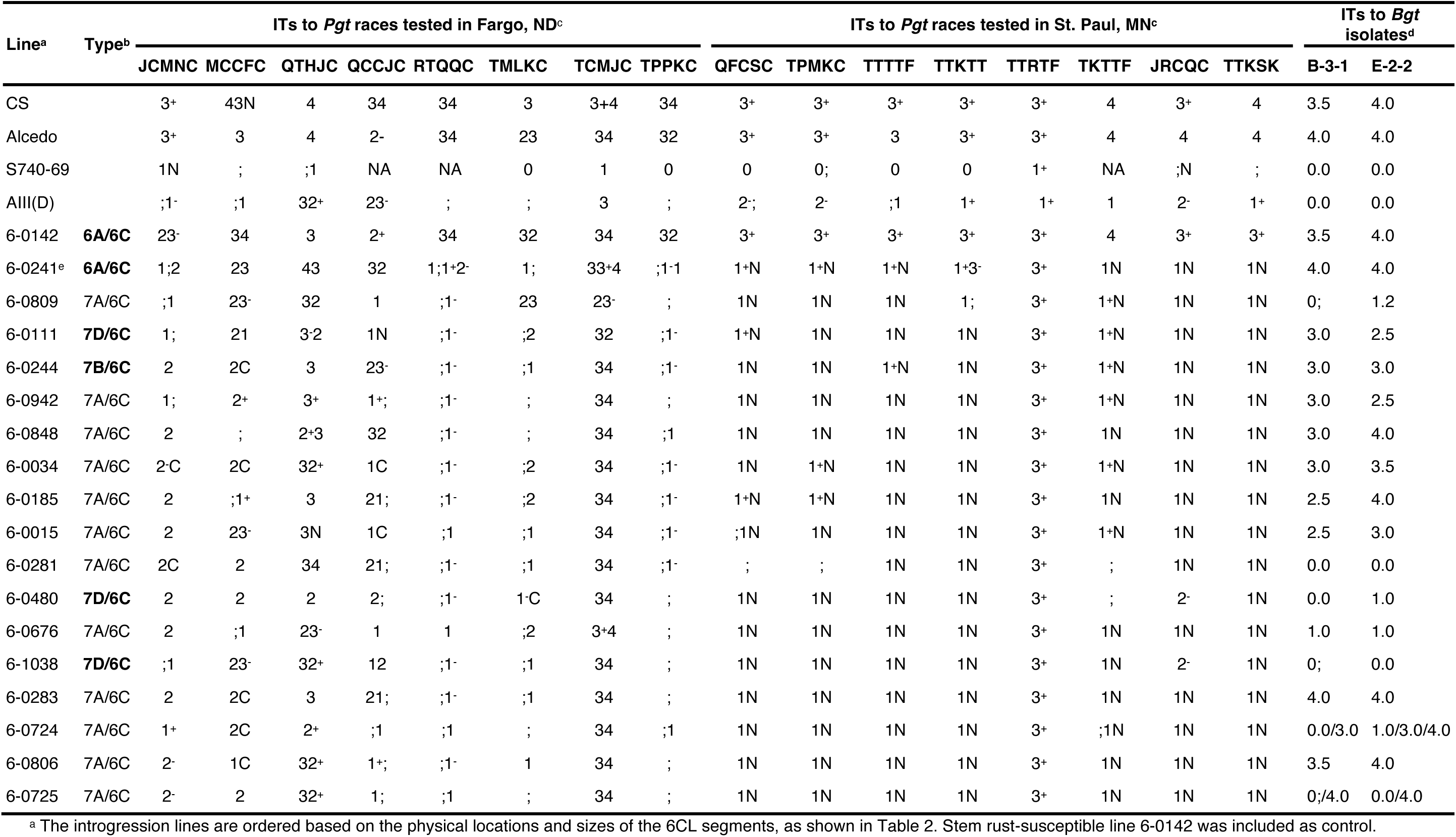
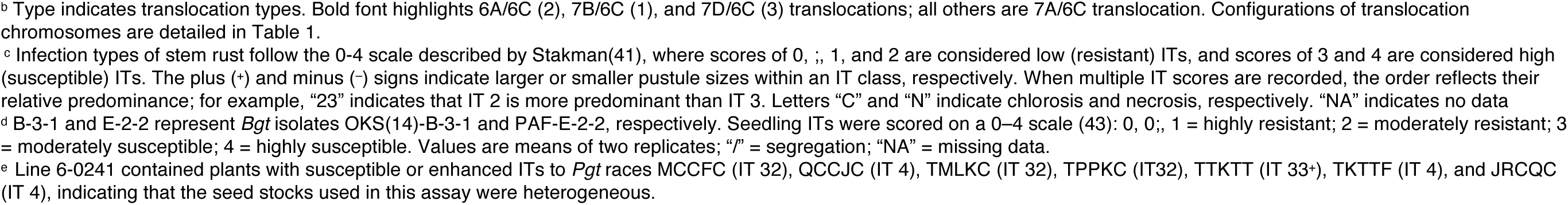
Infection types (ITs) of 17 Sr69 introgression lines and their parents including wheat Alcedo, the Ae. *caudata* S740-69 disomic addition line AIII(D), and Chinese Spring (CS) against 16 P. *graminis* f. sp. *tritici* (Pgt) races and two *Blumeria graminis* f. sp. *tritici* (*Bgt*) isolates tested in the greenhouses.

To assess temperature effects, three lines (6-0015, 6-0283, and 6-0480) were tested with eight races under greenhouse (20–23 °C) and growth chamber (18 °C and 25 °C) conditions (Fig. 2D and *SI Appendix*, Table S9). The three lines maintained their resistance spectrums across temperatures. However, temperature influenced infection response: at 25 °C, reactions showed increased necrosis, whereas at 18 °C, infection types tended toward mesothetic. Notably, responses to QTHJC were higher in growth chambers at both 18 °C (IT 231X) and 25 °C (ITs 12N–32) than under greenhouse conditions (IT 12).

Powdery mildew assays further revealed variation among introgression lines. Six lines (6-0281, 6-0480, 6-0676, 6-0724, 6-0809, and 6-1038) retained resistance from AIII(D), whereas the remaining 11 were susceptible (Fig. 2E and *SI Appendix*, Table S10 and S11). Because seed stocks from two to four BC₂F₃–BC₂F₄ families per line were tested, several lines segregated for mildew response. For example, lines 6-0724 and 6-0725—previously classified as resistant and susceptible, respectively, both segregated in greenhouse assays. Marker and GISH analyses showed that five resistant lines carried shorter introgressed 6CL segments than eight of the 11 susceptible lines (Fig. 1C), thereby preventing precise localization of the *Pm* gene(s). Because 6CL segment variation occurred mainly at proximal breakpoints, the *Pm* locus is likely located distal to *Sr69*. Small telomeric deletions or exchanges, which are difficult to detect by GISH and markers, may explain the loss of mildew resistance in susceptible lines.

## Discussion

Wild relatives are major sources of genetic variation for improving modern wheat but structural divergence between wild donors and wheat has long been considered a barrier to efficient and predictable recombination (26, 27). However, successful transfer of *Pm7C* (22), *Sr69*, and a potentially unmapped *Pm* gene from structurally rearranged *Ae. caudata* chromosomes demonstrate that extensive chromosomal divergence does not preclude precise, stable introgression. Instead, recombination outcomes appear to be governed primarily by local structural context and sequence similarity rather than classical homoeologous group relationships.

Here, we engineered a chromosome constitution containing a single wheat chromosome 6A and a single 6AS•6CL Robertsonian translocation to promote *ph1b*-mediated recombination between 6AL and 6CL. Unexpectedly, the rearranged 6CL segment preferentially recombined with its allosyntenic regions on chromosomes 7A, 7B, and 7D, despite these group-7 chromosomes being present as homologous pairs. This observation challenges the assumption that recombination in *ph1b* backgrounds primarily follows homoeologous group relationships and instead highlights the dominant role of chromosome-scale structural collinearity. The strong predominance of 7A/6C and 7D/6C translocations (16 lines), contrasted with the rarity of canonical 6A/6C recombination (one line), indicates that recombination can deviate substantially from expected homoeologous paring relationships.

Comparative genomic analyses provide a structural explanation for this pattern. The terminal ∼67 Mb of 6CL is syntenic with wheat group-7 telomeric regions rather than group-6 chromosomes (15), and recombination breakpoints are concentrated within this region. The close correspondence between rearranged chromosome structure and recombination outcomes indicates that recombination preferentially occurs within regions that share extended structural alignment, even when they reside on non-homoeologous chromosomes (Table 2). Together, these results provide clear evidence that *ph1b*-induced allosyndetic recombination can track syntenic regions among non-homoeologous chromosomes and can often be anticipated once chromosome-scale assemblies are available.

Sequence similarity analyses across genomic scales further clarify constraints on recombination partner choice. The approximately 4 percentage point increase in sequence similarity between 6CL and group-7 chromosomes, relative to group-6, likely represents a threshold that influences partner selection and the frequency of homoeologous recombination when *Ph1*-mediated constraints are removed (Fig. 2B). A similar pattern governs the *ph1b*-facilitated transfer of *Pm7C* from 7CL to 7DS (22), where sequence mismatches are accommodated at a rate commensurate with the overall divergence between chromosome pairs, conferring a recombination advantage on more closely related sequences (28). This principle likely also explains the preferential introgression of *Sr68* from a *Thinopyrum junceum* group-4 chromosome to wheat 1BS (11); such a transfer suggests the gene resides within a rearranged *Th. junceum* segment that is allosyntenic with 1BS.

Collectively, this study demonstrates that alien chromosomes with complex structures can be utilized predictably when their structural organization is well characterized, highlighting the importance of integrating structural genomics into chromosome engineering strategy. As genome assemblies for numerous wild relatives of wheat and other crops continue to expand, the framework established here expands access to previously underutilized diversity embedded in highly rearranged genomes. These findings support a genome structure–guided strategy for more efficient alien gene introgression for crop improvement.

## Materials and Methods

### Plant Materials

*Ae. caudata* accession S740-69, wheat cultivars ‘Alcedo’ and ‘Chinese Spring’ (CS), Alcedo-*Ae. caudata* S740-69 disomic addition line AIII(D), CS monosomic 6A (CS M6A), CS *ph1b* mutant, and CS nullisomic-tetrasomic lines (N3DT3B, N6DT6B, N7AT7B, N7AT7D, N7BT7A, N7BT7D, N7DT7A, and N7DT7B) were used in this study.

### Development of allosyndetic recombinants carrying stem rust resistance from *Ae. caudata*

A large BC₂F₁ population, which was used to select allosyndetic recombinants with minimal *Ae. caudata* chromatin carrying the *Sr* gene, was developed following previously described methods (11). Details regarding hybridization and marker-assisted selection for *ph1b* (29) are provided in the *SI Appendix*, *Method S1* and Fig. S1. The pedigree of the selection population was CS/4/CS *ph1b*/3/CS M6A/AIII(D)//CS *ph1b*.

### Molecular marker development and genotyping for idenfying wheat-*Ae. caudata* allosyndetic recombinants

Twenty-nine SSR markers polymorphic between CS and AIII(D) (21, 30–32) were initially screened on two stem rust-resistant and two susceptible BC₂F₁ plants to identify markers for allosyndetic recombinants. CS, AIII(D), and CS N3DT3B and CS N6DT6B served as controls. Selected markers (*SI Appendix*, Table S2) were used to genotype all resistant plants and a subset of susceptible BC₂F₁ plants by capillary electrophoresis (ABI 3130xl) (12). Additional SSR/STS markers (14 group-6, 20 group-7) were screened on 6% polyacrylamide gels, with PCR products stained with GelRed and imaged on a Typhoon 9410 scanner. Of these, only *BF145935* (*SI Appendix*, Table S3) was identified as co-dominant and used for characterizing translocation chromosomes in the allosyndetic recombinants.

STARP markers were developed following a previously described method (33). Low-copy CS sequences in regions replaced by *Ae. caudata* segments were identified via IWGSC BLAST and queried against *Ae. caudata* sequence read archive (SRA, SRX209239, SRX209237). Sequences with SNP/INDEL variation were used for allele-specific and common primers; reverse primers included ≥ 2 3′-end mismatches to reduce non-specific amplification. STARP markers were evaluated at 6.5% denaturing polyacrylamide gels (Li-Cor IR2 4300/4200) and validated with a CFX384 Real-Time PCR system (Bio-Rad). Twenty-two primer combinations were tested on CS, S740, AIII(D), and homozygous 7A/6C and 7D/6C lines. Validated co-dominant markers (*SI Appendix*, Table S3) were then used to genotype the BC₂F₂ populations of the allosyndetic recombinants.

### Cytogenetic analysis

Fluorescence GISH was performed following a previously described protocol (11). *Ae. caudata* genomic DNA labeled with biotin-16-dUTP (Enzo Life Sciences) served as probe, and sheared CS DNA was used as blocking DNA. Signals were detected with FITC-avidin (Vector Laboratories), and chromosomes were counterstained with propidium iodide (PI). For translocation identification, the same slides were used for oligo-FISH, employing TAMRA-modified oligos (pAs1-1, -3, -4, -6, AFA-3, -4) and FAM-modified oligos (pSc119.2-1, (GAA)₁₀) as probes. Chromosomes were counterstained with DAPI. Images were captured with an Axiocam HRm CCD on a Zeiss Axioplan 2 microscope and analyzed using AxioVision 4.5 (Carl Zeiss).

### Physical and Bionano optical mapping of *Ae. caudata* 6CL introgressions

Homoeolog-specific markers from syntenic genes were developed to identify *Ae. caudata* 6CL segments in wheat introgression lines. Proteomes of CS (IWGSC RefSeq v2.1) and *Ae. caudata* S740-69 (Aecau_v1) were compared using DIAMOND 2.1.8 (parameters: blastp --evalue 1e-9 --max-target-seqs 5) (34). Syntenic genes were identified using MCScanX v1.0.0 (35). PCR markers, spanning ∼60-Mb of 6CL region with ∼5 Mb intervals, were designed in Geneious Prime 2025.0, aligned with MUSCLE 5.1(36), targeting 100–200 bp genome-specific fragments with Tm ≈ 60 °C (Primer3 2.3.7). A 5′ sequence (GCAACAGGAACCAGCTATGAC) was appended for fragment size separation. PCR reactions (10 µL) contained 30–50 ng DNA, Taq buffer, dNTPs, primers (PEA1-700/800, forward, reverse), and 0.25 U Taq, cycled: 95 °C 3 min; 35× (94 °C 30 s, 60 °C 45 s, 72 °C 1 min); final extension 72 °C 7 min. Products were resolved on denaturing polyacrylamide gels (IR2 4300 DNA Analyzer, LI-COR).

Bionano optical mapping was used to validate introgression in line 6-0244 (Bionano Genomics Saphyr System). High-molecular-weight DNA was extracted from seedlings after three days dark treatment. Molecules were imaged on Saphyr Chip G2.3, exported as BNX files, and aligned to CS (IWGSC v2.1) and *Ae. caudata* Aecau_v1 using Bionano Solve 3.8.2 (*P* ≤ 1×10⁻¹⁰). Coverage depth was smoothed with a 1,000-site sliding window and visualized in R (v4.3.2) using ggplot2 and zoo (37, 38).

### Transmission analysis of introgressed segments in allosyndetic recombinants

Transmissions of *Ae. caudata* 6CL segments were evaluated in BC_2_F_2_ populations and BC_2_F_2:3_ families of 17 resistant recombinants (Table 2). For each line, 15–30 BC_2_F_2_ plants were inoculated with *Pgt* race TMLKC. Resistant BC_2_F_2_ plants were grown to produce BC_2_F_3_ seed, with ∼25–30 BC_2_F_3_ plants per BC_2_F_2_ plant tested with TMLKC. BC_2_F_2:3_ families segregating for resistance were first assessed for homogeneity using χ² tests. Heterogeneous families were sequentially removed until a homogeneous set remained, which was then tested for Mendelian segregation against the expected 3:1 ratio.

### Sequence similarity analysis

Pairwise alignments between the 6CL syntenic region and corresponding CS chromosomes (IWGSC RefSeq v2.1) (25) were performed with Minimap2 v2.28 (-f 0.0002) (39). Syntenic regions of group-7 chromosomes (7A, 7B, 7D) and size-matched terminal regions of group-6 chromosomes (6A, 6B, 6D) were aligned; for 7CL, alignments included wheat group-7 short arms (7AS, 7BS, 7DS) and matched long-arm regions as non-syntenic controls. Only alignment blocks with mapping quality = 60 were retained. Percentage identity was calculated as (matching residues/alignment length) × 100. Sliding-window analysis (1–200 adjacent blocks: 1, 10, 20, 30, 40, 100, 200) determined genomic span and mean identity. Pairwise comparisons among six chromosomes were assessed using Games-Howell’s test in Prism 10.2.3 (GraphPad Software, Boston, MA, USA).

### Evaluation of stem rust and powdery mildew resistance

Stem rust assays were performed using 16 *Pgt* races (*SI Appendix*, Table S1) at USDA-ARS, Northern Crops Science Laboratory (Fargo, ND, USA) and Cereal Disease Laboratory (St. Paul, MN, USA) following procedures previously described (11, 40, 41). Twenty-one *Bgt* isolates (*SI Appendix*, Table S10) collected in the U.S. were used to evaluate all wheat-*Ae. caudata* introgression lines for resistance to powdery mildew at two USDA-ARS locations (Raleigh, NC and Stillwater, OK). At Raleigh, tests were conducted using the detached-leaf method (21, 42) with 20 *Bgt* isolates. At Stillwater, lines were evaluated for responses to two *Bgt* isolates, OKS(14)-B-3-1 and PAF-E-2-2, in greenhouse using the protocol previously described (43, 44).

## Acknowledgments

The authors thank Shiaoman Chao and Mary Osenga for assistance with technical support in capillary electrophoresis, Danielle Holmes for technical support in stem rust testing, and Han-Chang Chang for seed increase. This project is a collaborative effort established through Non-Assistance Cooperative Agreements between USDA-ARS and the University of California at Davis (Agreement No. 58-2030-4-085), the University of California at Berkeley (Agreement No. 58-2030-2-032), and North Dakota State University (Agreement No. 58-3060-9-031). This research was supported in part by funds to S. S. X. provided through a grant from the Bill & Melinda Gates Foundation and UK Department for International Development to Cornell University for the Borlaug Global Rust Initiative (BGRI) Durable Rust Resistance in Wheat (DRRW) Project (Grant Number OPPGD1389) and the USDA-ARS CRIS Project No. 3060-21000-046-000-D and 2030-21430-015-000-D. Mention of trade names or commercial products in this article is solely for the purpose of providing specific information and does not imply recommendation or endorsement by the USDA.

## Author Contributions

S.S.X. and D.L.K. initiated and conceived the study. H.-L.Y., D.L.K., Q.Z., Z.N. Y.Q.G., X.Z., Y.J., S.G., G.Q.L., R.B.W., A.P.H., T.L.F., X.Y.X., S.M.Y., C.C., J.D.F., S.L., and X.Y.C., conducted research; H.-L.Y., S.N.M., X.F.Z., D.L.K., P.C., J.S., J.L.C., and S.S.X. analyzed the data; S.S.X. and Y.Q.G. acquired funding; H.-L.Y., S.N.M., X.F.Z., D.L.K. and S.S.X. drafted the manuscript. All authors reviewed and revised the manuscript.

## Competing Interest Statement

The authors declare no conflicts of interest.

**Additional Supporting Data and Tables**

**Supplemental Figures**

## Supplementary Information Appendix

**This file includes:**

*SI Appendix*, Method S1

*SI Appendix*, Figures S1 to S9

*SI Appendix, References*

### *SI Appendix*, Methods

#### SI Appendix, Method S1. Production of Robertsonian translocations and allosyndetic recombinants

Allosyndetic recombinants carrying stem rust resistance gene *Sr69* from wheat ‘Alcedo’-*Aegilops caudata* S740-69 disomic addition line AIII(D) were developed using the method as previously described (1). To generate a Robertsonian translocation, ‘Chinese Spring’ (CS) monosomic 6A (CS M6A) was crossed as the female parent with AIII(D) (SI Appendix, Fig. S1). The F_2_ plants derived from double monosomic F₁ plants (2*n* = 42, 20″ + 1′_6A_ + 1′_6C_) were evaluated for reaction to *Pgt* race TMLKC and resistant plants with infection types (IT) ≤ 23 were subjected to genomic *in situ* hybridization (GISH) analysis following Mandel et al (1) except that root tips were collected after stem rust phenotyping. An F_2_ plant (CS M6A/AIII(D)) carrying a 6AS•6CL Robertsonian translocation chromosome was used as the female parent in a cross with the CS *ph1b* mutant. The resulting F₁ plants were evaluated with *Pgt* race TMLKC, and resistant plants were backcrossed to the CS *ph1b* mutant. BC₁F₁ plants were again tested for stem rust resistance using race TMLKC. Leaf tissues were collected from resistant BC₁F₁ plants for DNA extraction as described below. Samples were analyzed using multiplex PCR with markers *AWJL3*, *Psr574*, and *Psr128*. *AWJL3* served as a positive control, whereas *Psr574* and *Psr128* were used to detect the absence of the 5BL segment carrying *Ph1* (2). Rust-resistant BC_1_F_1_ plants lacking *Ph1* were crossed as male to CS to generate a large BC_2_F_1_ population for selection of allosyndetic recombinants carrying shortened alien chromatin segments. The pedigree of the selection population was CS/4/CS *ph1b*/3/CS M6A 6A/AIII(D)//CS *ph1b*. The allosyndetic recombinants carrying *Sr69* on the short *Ae. caudata* chromosome segments were identified from the large BC_2_F_1_ population based on stem rust test, marker genotyping, and GISH analysis.

### *SI Appendix*, Figures

**Fig. S1.**
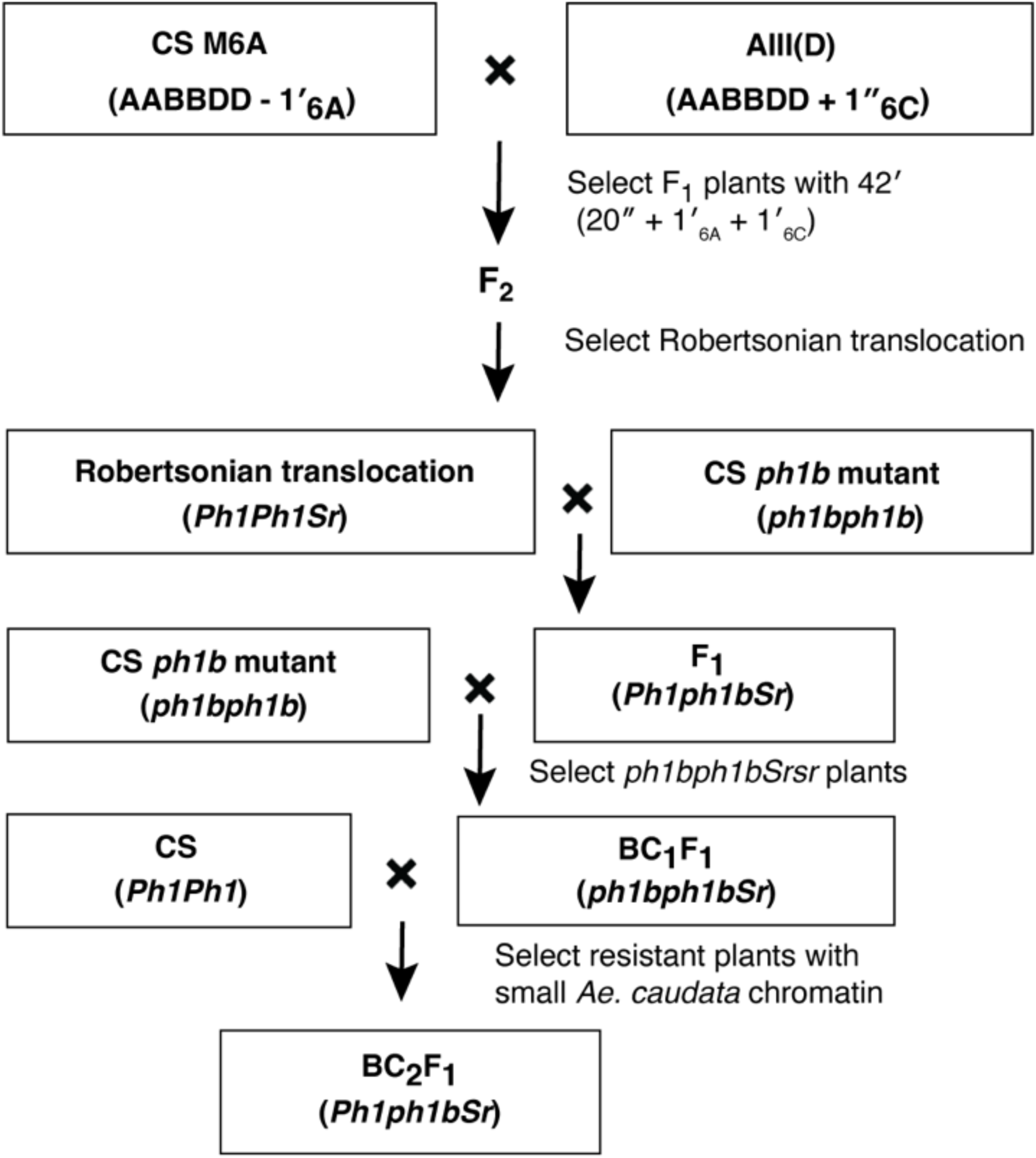
Hybridization and selection scheme for generating allosyndetic recombinants of hexaploid wheat carrying stem rust gene *Sr69* and powdery mildew resistance gene(s). CS M6A refers to the ‘Chinese Spring’ (CS) monosomic 6A line; AIII(D) is the disomic addition line carrying a pair of 6C chromosomes from *Ae. caudata* accession S740-69 in the ‘Alcedo’ wheat background.

**Fig. S2.**
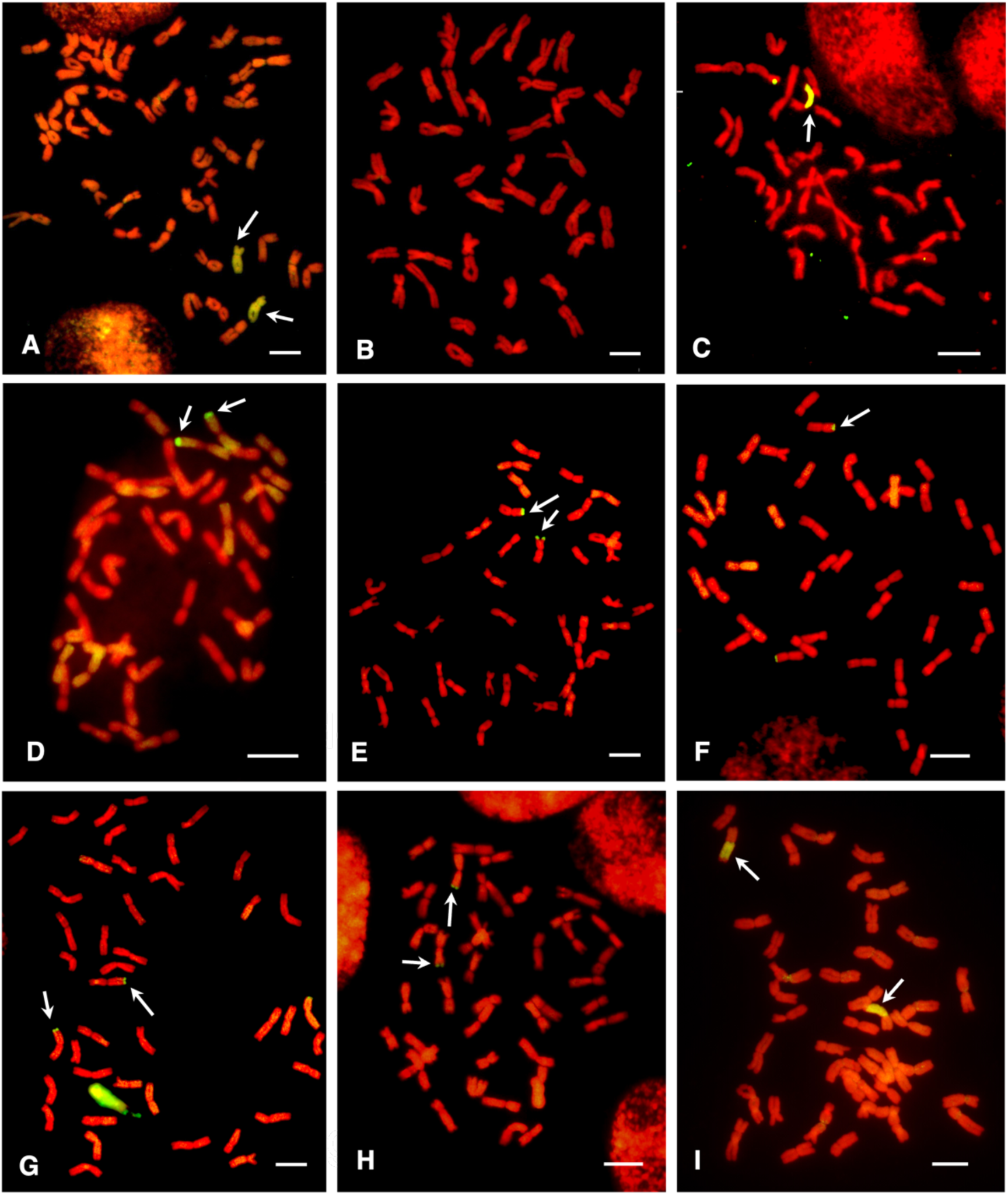
Genomic *in situ* hybridization analysis of wheat-*Ae. caudata* introgression lines carrying stem rust resistance gene *Sr69*. *Ae. caudata* chromatin (arrows) was labeled with fluorescein isothiocyanate–conjugated avidin (green). **A.** ‘Alcedo’-*Ae. caudata* S740-69 disomic addition line AIII(D). **B.** Chinese Spring (CS). **C.** Original 6AS•6CL Robertsonian translocation plant (12N61-16). **D.** 6-0809 (6A/6C recombinant). **E.** 6-0111 (7D/6C). **F.** 6-0480 (7D/6C). **G.** 6-0015 (7A/6C). **H.** 6-0281 (7A/7C), and **I.** 6-0303 (6A/6C). The introgressed segments carrying *Sr69* are telomeric Conversely, in the susceptible line 6-0303 (**I**), the distal *Ae. caudata* segment has been replaced by wheat chromatin. Scale bar = 10 µm.

**Fig. S3.**
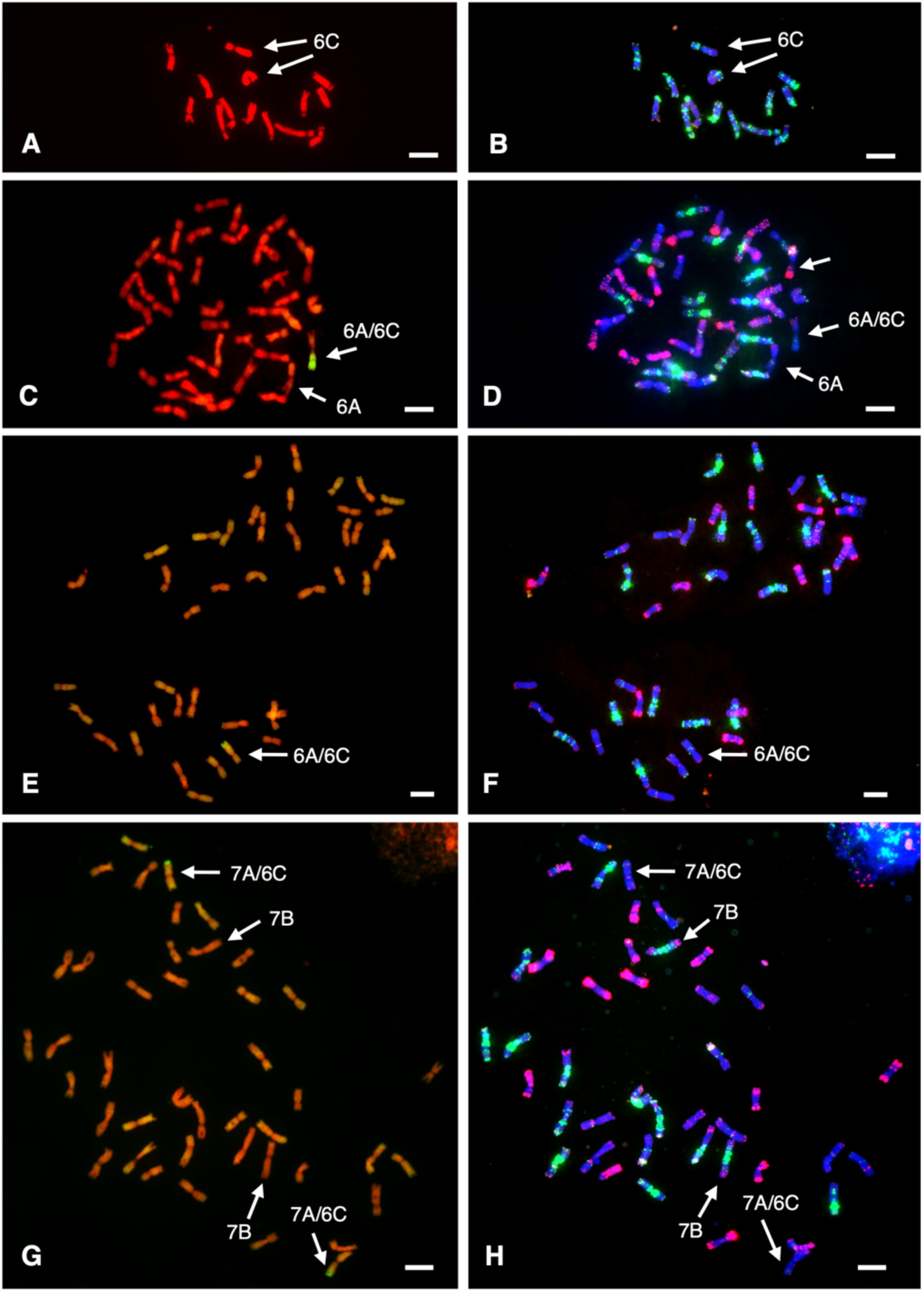
Identification of translocation chromosomes in wheat-*Ae. caudata* introgression lines 6-0142, 6-0241, and 6-0034 (control) using GISH and oligo-FISH. Each line is shown with GISH (left) and oligo-FISH (right) images, except *Ae. caudata* S740-69 (control), which shows chromosomes stained with DAPI (**A**) prior to oligo-FISH analysis (**B**). Lines 6-0142 (**C, D**) and 6-0241 (**E, F**) carry 6A/6C translocations, whereas 6-0034 (**G, H**) carries a 7A/6C translocation. Scale bar = 10 µm.

**Fig. S4.**
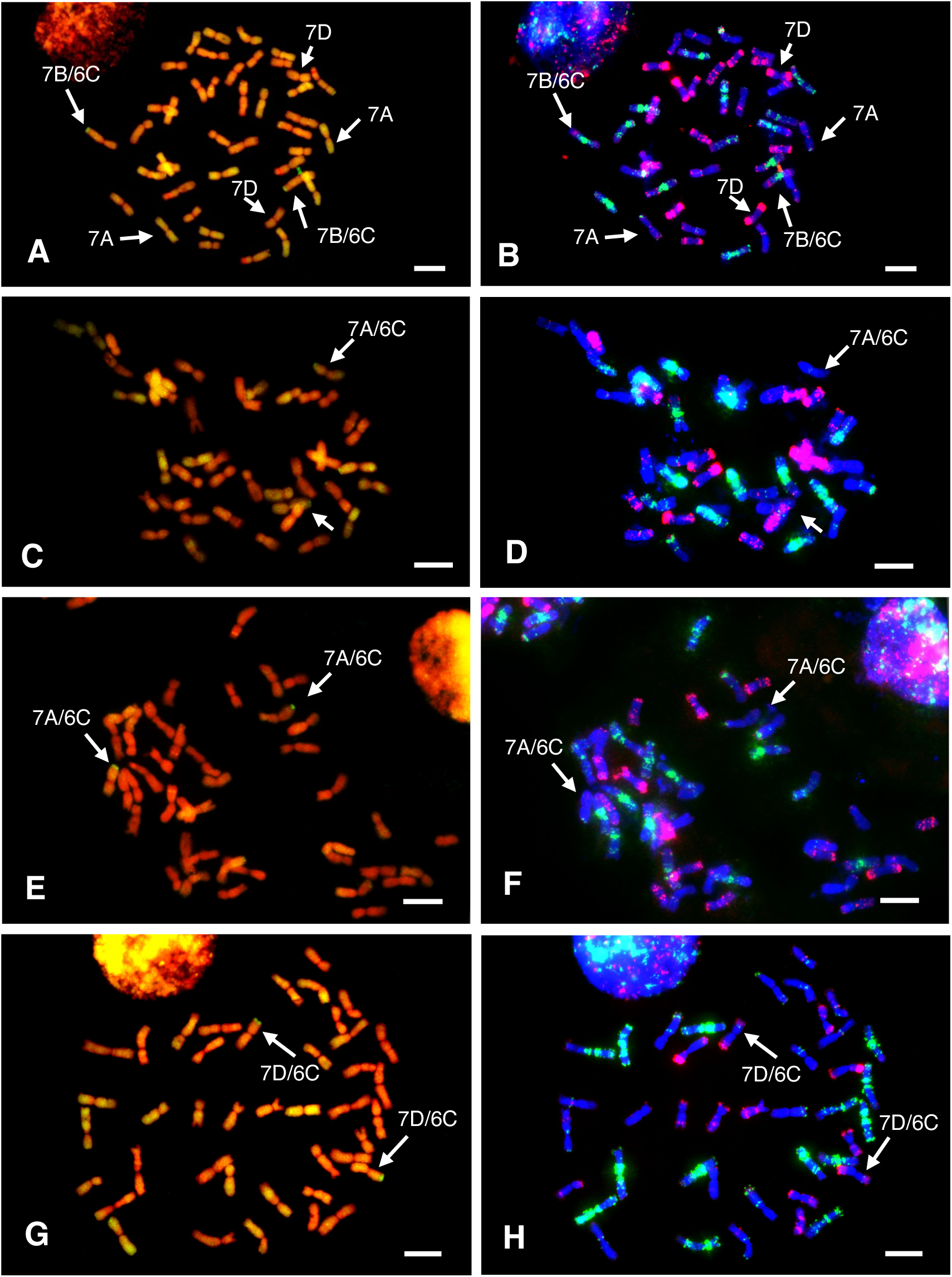
Identification of translocation chromosomes in wheat-*Aegilops caudata* introgression lines, 6-0244, 6-0725, and 6-0806, using GISH and oligo-FISH. Each line has two images from GISH (left) and oligo-FISH (right) analysis. Line 6-0244 (**A, B**) carries a 7B/6C translocation, whereas 6-0725 (**C, D**) and 6-0806 carry 7A/6C (**E, F**) translocations. Line 1038 carries a 7D/6C translocation (**G, H**). Bar = 10 µm.

**Fig. S5.**
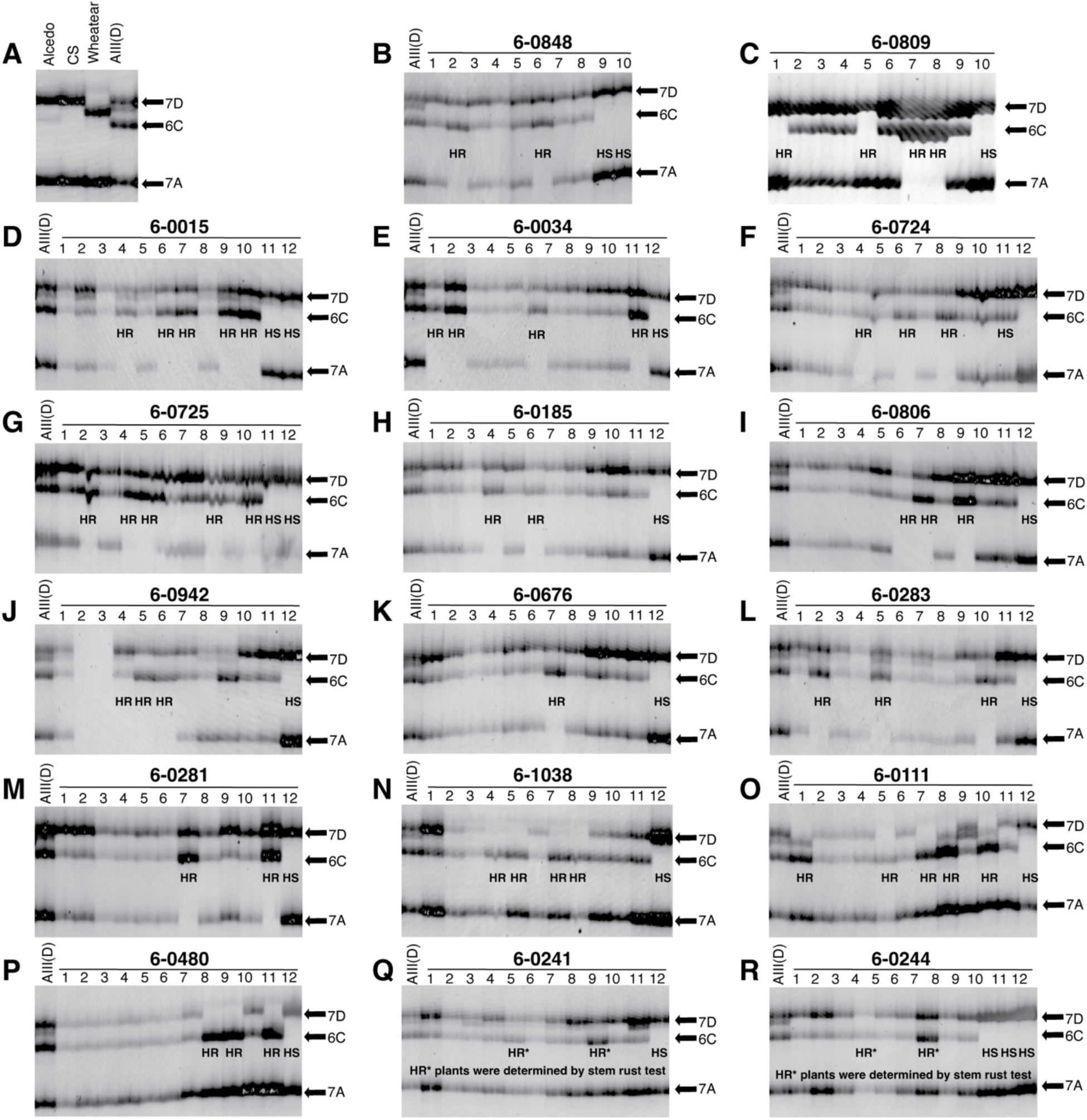
Analysis of BC₂F₂ populations from 17 wheat–*Ae. caudata* recombinant (BC₂F₁) plants using marker *BF145935*. The numbers 1–12 below each line indicate individual BC₂F₂ plants. Homozygous stem rust–resistant (HR) and susceptible (HS) plants were confirmed by stem rust testing. **A.** Gel image showing 7A, 7D, and 6C bands in wheat parents (‘Alcedo’, CS, AIII(D), and Wheatear) based on nullisomic–tetrasomic analysis (Fig. 1A). **B–M.** Twelve lines carried 7A/7C translocations, indicated by the absence of the 7A band in HR plants. **N–P**. Three lines carried 7D/7C translocations, indicated by the absence of the 7D band in HR families. **Q–R.** Two lines (6-0241 and 6-0244) carried translocations that could not be determined using marker *BF145935*.

**Fig. S6.**
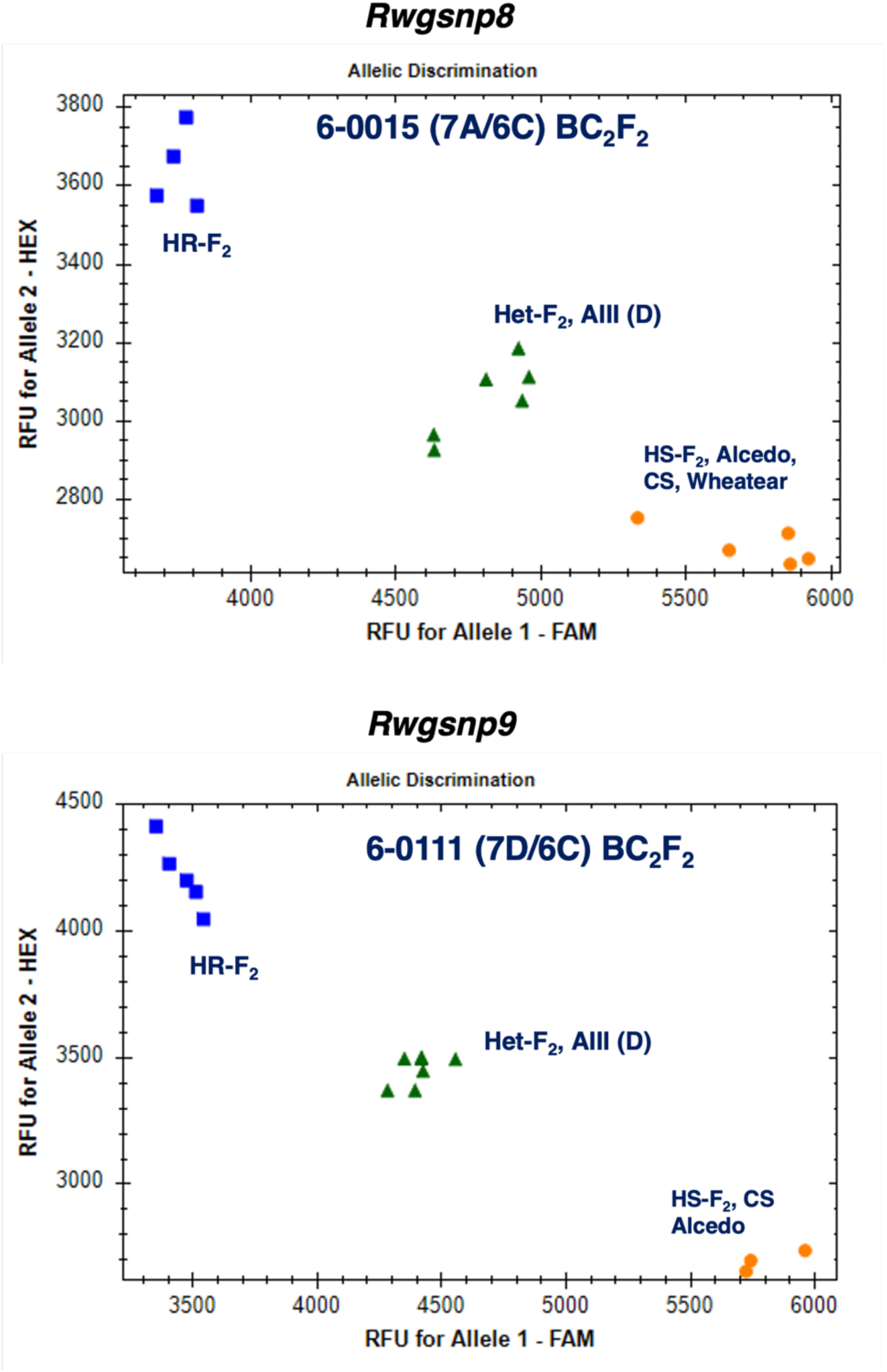
Analysis of BC₂F₂ populations of 6-0015 (7A/6C) and 6-0111 (7D/6C) using STARP markers Rwgsnp8 and Rwgsnp9 with the CFX384 Touch™ Real-Time PCR Detection System. Each population was phenotyped with *Puccinia graminis* f. sp. *tritic*i race TMLKC prior to DNA extraction. In the genotype clusters, HR, Het, and HS indicate homozygous stem rust–resistant, heterozygous, and susceptible BC₂F₂ plants, respectively (see Fig. 1B).

**Fig. S7.**
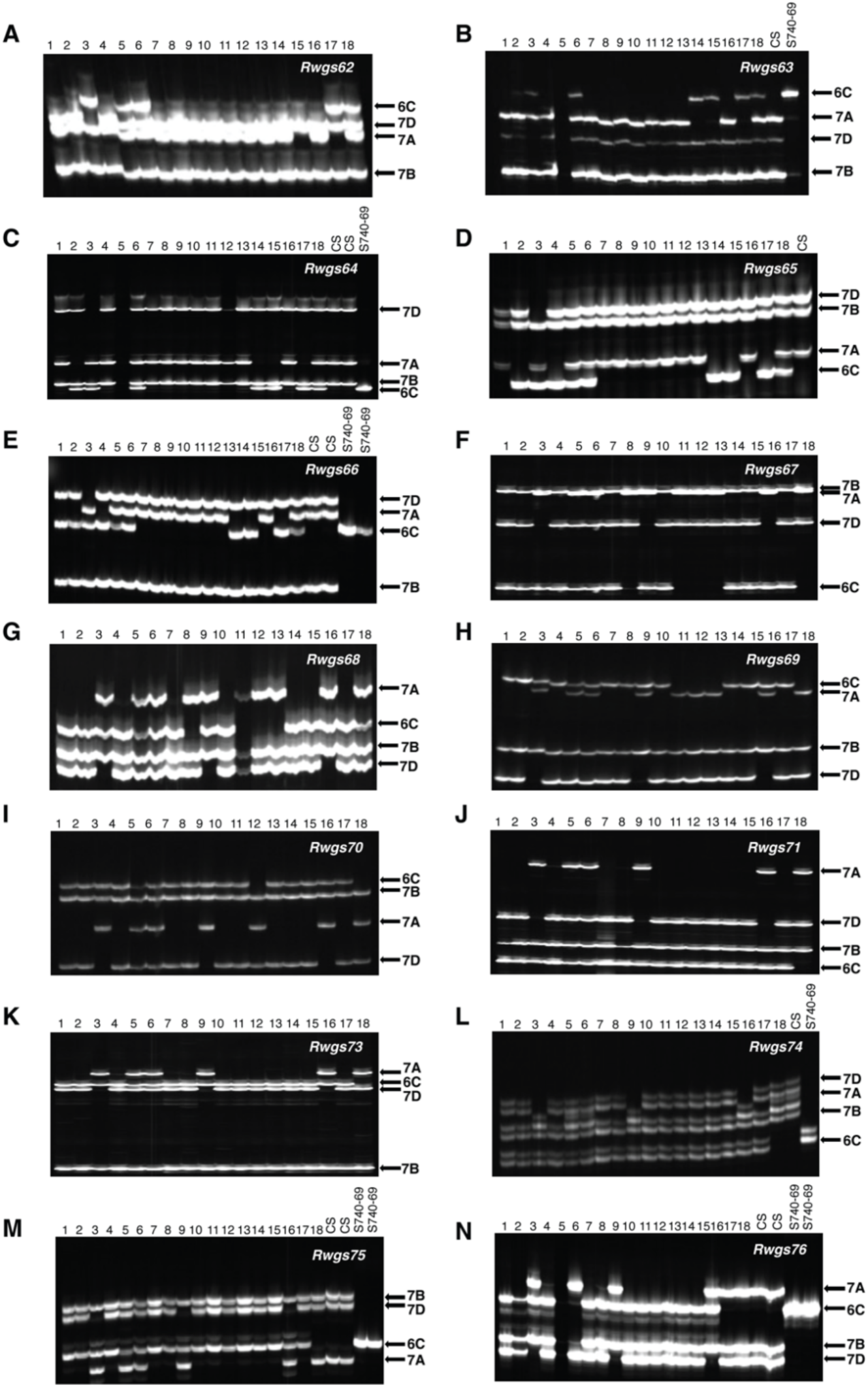
Gel images of 14 syntenic gene-specific markers from genotyping 17 *Sr69* introgression lines and controls. The numbers on the top of each gel image represent introgression lines and controls: 1 = 6-0015, 2 = 6-0034, 3 = 6-0111, 4 = 6-0185, 5 = 6-0241, 6 = 6-0244, 7 = 6-0281, 8 = 6-0283, 9 = 6-0480, 10 = 6-0676, 11 = 6-0724, 12 = 6-0725, 13 = 6-0806, 14 = 6-0848, 15 = 6-0942, 16 = 6-1038, 17 = 6-0809, 18 = 6-0142, CS = Chinese Spring, S740-69 = *Ae. caudata* S740-69. Wheat chromosomes involving in the translocation in three lines, 6-0142, 6-0241 and 6-0244, could not be determined by all these markers.

**Fig. S8.**
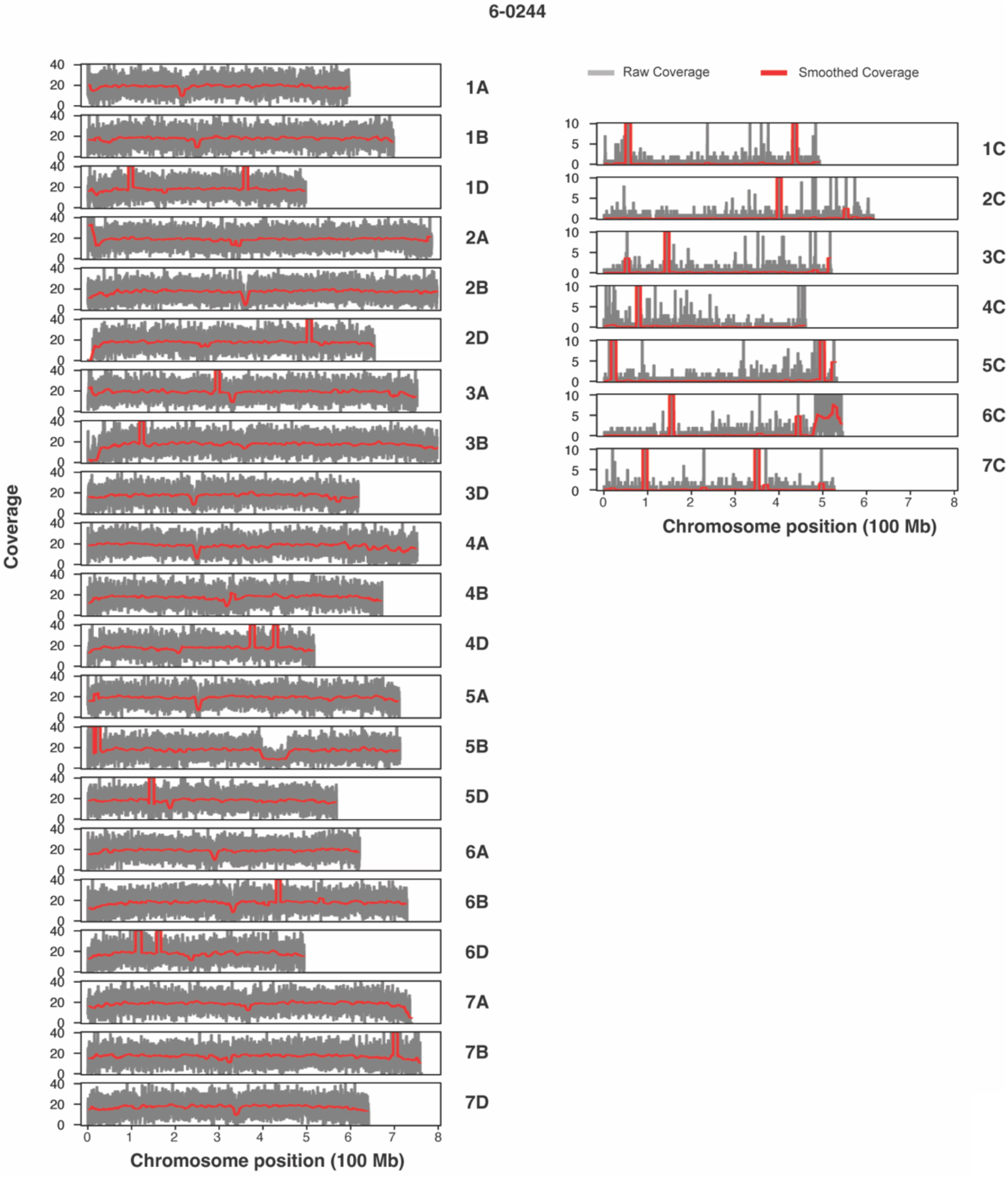
Bionano optical mapping of wheat-*Ae. caudata* introgression line 6-0244 showing alignment coverage against *Ae. caudata* Aecau_v1 (3) and CS IWGSC RefSeq v2.1 (4) genomes. A 59.3-Mb *Ae. caudata* segment was mapped to the distal end (483.15–542.49 Mb) of chromosome arm 6CL, a 5-Mb telomeric regions were missing on wheat chromosome arms 7AL, whereas no significant deletion was detected on chromosome 7B.

**Fig. S9.**
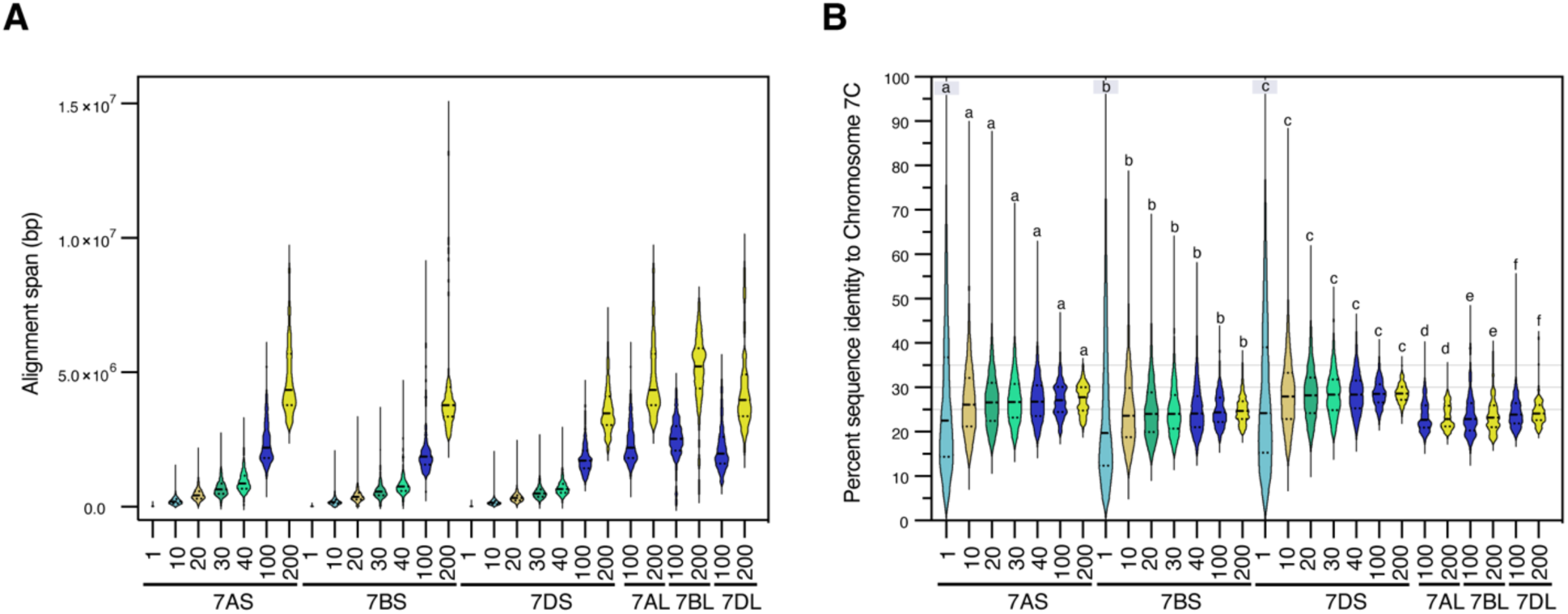
Alignment between the *Ae. caudata* 6CL( Left) and 7CL (Right) syntenic region and each corresponding group-7 chromosome segment. **(A)** Pairwise alignment span (bp) of the 6CL syntenic region against whole chromosomes 7A, 7B, 7D, and group-6 chromosomes 6A, 6B, and 6D. **(B)** Pairwise alignment span of the 7CL syntenic region against the short arms (7AS, 7BS, 7DS) and long arms (7AL, 7BL, 7DL) of group-7 chromosomes.

